# Microencapsulation of Nisin in Polyelectric complexes of alginate-chitosan for extended antimicrobial activity

**DOI:** 10.64898/2026.08.14.744846

**Authors:** Christian Kosisochukwu Anumudu, Taghi Miri, Helen Onyeaka

## Abstract

Nisin is a promising antimicrobial peptide widely used in food preservation due to its efficacy against Gram-positive spoilage and pathogenic bacteria. Although Nisin is increasingly applied in the food sector, the biopeptide suffers from instability within food matrixes and can rapidly lose its antimicrobial potential following interaction with food biomolecules. Thus, it is necessary to investigate approaches that can be employed to extend the stability and activity of Nisin. Hence, the aim of this study was to develop and characterise a chitosan–alginate polyelectrolyte microencapsulation system capable of enhancing Nisin stability while retaining antimicrobial activity. The microencapsulation of Nisin was achieved by pre-gelation of alginate using calcium chloride and subsequent direct electrostatic interaction between cationic Nisin and chitosan with pre-gelled anionic alginate at pH 5.0. Following microcapsule formation, physicochemical and structural characterisation was performed using Zeta potential determination and measurement of the polydispersity index (PDI) via dynamic light scattering. SEM micrographs were used to confirm morphology, while Fourier-transform infrared (FTIR) spectroscopy and high-performance liquid chromatography (HPLC) were utilised to assess chemical integrity and functional group preservation of encapsulated Nisin. Following this, stable microcapsules with diameters ranging from 150–200 nm and smooth surface morphology were obtained. Microcapsule formation was strongly influenced by formulation parameters, particularly pH, calcium ion concentration, and chitosan content, with deviations from optimal acidic conditions (< pH 5.0) resulting in aggregation, increased polydispersity, and reduced encapsulation efficiency. The microcapsules were monodispersed (PDI ≈ 0.30) and electrostatically stable, exhibiting a Zeta potential of approximately +36 mV. These microcapsules remained stable over a prolonged storage period of 21 days under refrigerated conditions while retaining antimicrobial activity against *Bacillus cereus*. Encapsulation efficiency reached approximately 65%, confirming effective retention of Nisin within the polymer matrix. Overall, the findings demonstrate that chitosan–alginate ionic gelation is a non-denaturing and effective encapsulation strategy for extending the functional stability of Nisin. These microcapsules show strong potential as natural antimicrobial delivery systems for food and beverage applications, particularly in acidic food matrices, with implications for improved food safety and shelf-life extension.

## 1. INTRODUCTION

Nisin has been applied widely in different food matrixes. The bacteriocin is active against a broad range of bacteria, including the gram-positive *Bacillus cereus.* Its activity against *Bacillus cereus* is important due to the burden of food spoilage and possible food-borne illnesses posed by the bacteria, especially in canned foods, due to its ability to produce endospores which are highly resistant to adverse conditions found in many processed foods which ordinarily would kill off the vegetative forms of many contaminating microorganisms. Furthermore, *Bacillus cereus’* survival in food poses a higher risk as the organism can produce enterotoxins, which has far-reaching consequences. Consequently, the use of nisin as a biopreservative represents an important strategy for improving the microbiological safety of processed and ready-to-eat foods. However, because of the peptide nature of Nisin, it is usually degraded rapidly within food matrixes, primarily upon heating at low pH. This degradation can be associated with enzymatic degradation [1, 2] and due to interaction with lipids and proteins within the foods. Thus, several approaches have been investigated to microencapsulate Nisin, shielding it from degradation within food matrixes and ensuring slow and continuous release of the peptides into the foods. Some of these approaches include the use of nano-liposomes [3], the use of chitosan alone [4], and alginate beads alone [5]. Despite these advances, limitations such as low encapsulation efficiency, instability during processing, and uncontrolled release kinetics remain challenges for the practical application of encapsulated nisin in food systems. The use of biopolymers for the delivery of antimicrobials is receiving increased attention, and this is attributable to the ease of fabrication, their biodegradability, digestibility and inexpensive nature [6, 7].

Chitosan is a positively charged carbohydrate polymer derived from the exoskeletons of many crustaceans. It is composed of 2-acetamido-2-deoxy-β-D-glucose units linked together by (1–4) glycosidic bonds, whereas alginate is an anionic polysaccharide copolymer of β-d-mannuronic acid (M) and α-L-guluronic acid (G) residues linked together by (1,4) glycosidic bonds [8]. Chitosan and alginate are utilised in the formation of microcapsules for drug delivery due to their inherent linkages and folding ability, properties which make them excellent candidates as delivery vehicles, especially for amphipathic Nisin. Nisin-alginate-chitosan microcapsules are formulated and stabilised as a result of the electrostatic interactions between positively charged Nisin and the negatively charged encapsulating biopolymers. Nisin-chitosan-alginate encapsulation is achieved through complexation in the presence of calcium ion (Ca^2+^). Under continuous stirring, alginate and calcium forms a stable calcium-alginate pre-gel having a negatively charged carboxyl groups presented, which can grow into a continuous polymeric network. When this is complexed with the positively charged amine groups of chitosan, it results in a stable cross-linked hydrogel which can be utilised to encase Nisin. This natural non-specific interaction between the charged molecules enables the rapid encapsulation of Nisin and can be utilised as an effective delivery system for the slow controlled release of Nisin compared to using either of the polymers alone. Previous research [9, 10] has studied these microcapsules as potential delivery systems for encapsulating several biomaterials, including bovine serum albumin (BSA), by alternating the ratio of chitosan to alginate to obtain enhanced entrapment. Usually, entrapment/encapsulation efficiency varies based on pH and polymer concentration for chitosan:alginate ratios of 1:1. An earlier study [11] which utilised an alginate-chitosan-pluronic composite for the encapsulation of Nisin obtained an encapsulation efficiency of 41.45% to 88.36% depending on formulation factors, with an average microcapsule diameter of 130–178 nm. Overall, several factors contribute to the success of Nisin encapsulation using chitosan:alginate, and these include pH, concentration of Ca^2+^ ions, agitation and concentration [8].

However, despite their inherent stability, current fabrication approaches for nisin–alginate–chitosan microcapsules often exhibit variable encapsulation efficiency and structural decoupling, limiting their effectiveness for extended antimicrobial activity [12]. In this study, a nisin–alginate–chitosan polyelectrolyte complex stabilised with D-trehalose was developed to improve encapsulation efficiency and achieve sustained antimicrobial release. The encapsulation efficiency, microcapsule morphology, particle size distribution, and zeta potential were evaluated, alongside the effect of varying calcium ion concentrations on microcapsule stability and physicochemical properties.

## 2. METHODOLOGY

### 2.1. Materials

Ultrapure Nisin with purity ≥99% (≥38,000IU/mg) was supplied by Handary S.A (Brussels, Belgium), chitosan (0.5%; LMW: 52 kDa) and 1% acetic acid was purchased from Sigma, USA and used to solubilise the chitosan. Sodium alginate (33% mannuronate and 67% guluronate) was purchased from Sigma-Aldrich (Saint Louis, USA). Muller Hinton broth (MHB), Tryptic Soy Broth (TSB) and Brain Heart Infusion Broth and Agar were purchased from Fisher Scientific (United Kingdom) Anhydrous calcium chloride and d-trehalose were obtained from ThermoFisher Scientific, UK. For chromatographic analysis, acetonitrile (99.99% purity) was purchased from Macron Chemicals (Pennsylvania, USA), and methanol (99.99% purity) was supplied by Fisher Scientific (Leicestershire, UK). Formic acid (98% purity) and acetic acid (analytical grade) was supplied by Fluka (Sigma-Aldrich, Missouri, USA). Hydrophilic 0.22 μm Millipore syringe filters were bought from Sigma-Aldrich, UK. Phosphate buffered saline (PBS) was prepared by dissolving commercial PBS granules purchased from Thermo Scientific (Rockford, USA) in ultrapure water. Ultrapure water was generated with a Milli-Q Purification System (Merck Millipore, USA). Calcium ion strength during encapsulation was monitored using a NexSens WQ-CA ISE Sensor (USA). A NE-1010 Syringe Pump (USA) was used for controlled titration of calcium chloride, while viscosity measurements were obtained using a Canon-Fenske Capillary Viscometer (Canon Instruments Corporation, USA). Particle size, polydispersity index, and Zeta potential analyses were conducted with a Zetasizer Pro (Malvern Instruments Ltd., UK). For chromatographic analysis, a Shimadzu Prominence HPLC system (Japan) equipped with a UV–Photodiode Array Detector and a C18 column (Thermo Fisher Scientific, UK) was used. Scanning electron microscopy (SEM) was performed with a Zeiss EVO 15 VP E-SEM (Zeiss Instruments, Germany), with sample preparation achieved using a Labconco Benchtop Freeze Dryer (Cole Parmer, UK) and gold sputter coating (Thermo Fisher Scientific, UK). FTIR spectra were collected using a Bruker IFS 66/s spectrometer (Germany) equipped with a Harrick MVP-Pro™ diamond ATR accessory. All other reagents were of analytical standard. *B. cereus* NCTC 11143 was supplied by the Biochemical Engineering Laboratory of the University of Birmingham.

### 2.2. Preparation of chitosan-alginate microcapsules

The preparation of the Nisin-alginate-chitosan was based on previously developed methods in the literature [8, 13] with modifications. This was achieved by an initial direct electrostatic interaction between cationic Nisin and chitosan with anionic alginate. Sodium alginate solutions (0.05%) were prepared by dissolving in distilled water. Similarly, chitosan (0.05%) was dissolved in 1% acetic acid with continuous stirring. Stock Nisin solution was prepared by dissolving in 0.05% acetic acid to a concentration of 1mg/ml. Solutions of calcium chloride (0-2 mM) were prepared by dissolving in distilled water. Calcium ion strength was measured using a NexSens WQ-CA ISE Sensor. Calcium chloride was titrated dropwise into the sodium alginate solution (ratio of 1:10) at a rate of 1 ml/min at 60 °C, which was selected to promote polymer mobility and complex formation while maintaining Nisin stability, using an NE 1010 syringe pump with continuous stirring (400 rpm) for 30 minutes to produce a calcium-alginate pre-gel. The viscosity of this calcium-alginate solution was measured with the aid of a Canon-Fenske capillary viscometer (Canon Instruments Corporation, USA) at 20□C following standard viscometric protocols to obtain the kinematic viscosity using the formula as outlined in Equation 1.

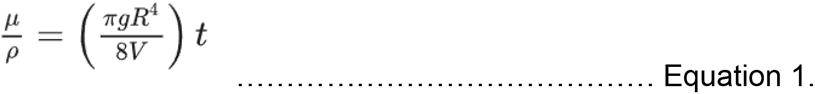

Where μ = solution viscosity (Pa.s), ρ = solution density (g/cm^3^), g = acceleration due to gravity (m/s^2^), R = capillary radius (m), V = volume of liquid (ml) and t = efflux time (s). The left-hand side of equation 1 represent measured variables whilst the right-hand side represent the capillary viscometer constant which has a value of 1.2 mm^2^/s^2^. The density of the solution μ was used to work out the relative viscosity (μr) using the viscosity of the solvent (0.05% sodium alginate in distilled water). This is outlined in Equation 2.

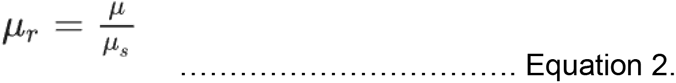

Where, μ is the solution viscosity, μs is the viscosity of the solvent (0.05% sodium alginate in distilled water) and μr is the relative viscosity.

Following this, Nisin solution was added dropwise into the sodium alginate pre-gel. Then, chitosan solution was precipitated into the calcium-alginate-Nisin solution at a 1 ml/min rate at 60 °C with continuous stirring (400 rpm). All solutions were mixed using a stir bar for 30 min (400 rpm) to ensure a homogenous mixture and complex formation at 60 °C. Following complex formation, the microcapsules were harvested by separation via centrifugation at an increased speed of 2000rpm for 30 min. Obtained particles were washed twice in PBS and frozen at −80 °C for 4 hours prior to further analyses.

### 2.3. Response Surface Methodology

The choice of optimal encapsulation approach to be utilized in this study was guided by statistical analysis based on response surface methodology (RSM), which was used to examine the effect of three independent formulation variables on microcapsule performance: chitosan concentration (mg/mL), the relative viscosity of the calcium-alginate-Nisin solution, and the CS:Nisin-alginate ratio.

A three-factor, fifteen-run experimental matrix was constructed, and data collected from each run were organised into a structured dataset (Table 1). Before fitting the models, the CS:Nisin-alginate ratio, which was initially categorical (1:1, 1:2, 2:1), was encoded numerically to allow for regression analysis. The relative viscosity and CS concentration were treated as continuous variables.

**Table 1.** Experimental response variables for response surface design.

| Experimental Run | Variables |  |  |
| --- | --- | --- | --- |
|  | CS Concentration (mg/mL) | Relative Viscosity | CS/Nisin-alginate Ratio |
| 1 | 0.5 | 0.8 | 1:1 |
| 2 | 0.5 | 1.2 | 1:1 |
| 3 | 0.5 | 1.0 | 1:1 |
| 4 | 1.0 | 0.8 | 1:2 |
| 5 | 1.0 | 1.2 | 1:2 |
| 6 | 1.0 | 1.0 | 1:2 |
| 7 | 1.5 | 0.8 | 2:1 |
| 8 | 1.5 | 1.2 | 2:1 |
| 9 | 1.5 | 1.0 | 2:1 |
| 10 | 1.0 | 1.0 | 1:2 |
| 11 | 0.75 | 1.0 | 1:2 |
| 12 | 1.25 | 1.0 | 1:2 |
| 13 | 1.0 | 0.9 | 1:2 |
| 14 | 1.0 | 1.1 | 1:2 |
| 15 | 1.0 | 1.0 | 1:1 |

Quadratic regression models were then fitted to the data using the ordinary least squares method in R (version 4.3.3), employing the LM and RSM packages. Each response variable (encapsulation rate, Zeta potential, and PDI) was modelled separately, including linear, quadratic, and two-factor interaction terms. Analysis of variance (ANOVA) was used to assess the statistical significance of each model term, with p-values less than 0.05 considered significant. The composite response variable (M) was defined as the sum of encapsulation efficiency and Zeta potential, serving as a practical indicator of overall performance.

To support interpreting the results and finding the best conditions, response surface plots were produced based on the fitted models. The ranges of the experimental variables were put in a standard form so their interactions could be easily seen. The diagrams showed which values for the input variables created the highest amount of the composite matter (M).

This method was chosen because RSM is effective for spotting unusual relationships and managing multivariate processes using a small amount of information. Using regression modelling, visual analysis and practical requirements, the statistical analysis built a reliable and easy-to-interpret framework for the formulation.

### 2.4. Evaluation of the stability of Nisin-chitosan-alginate microcapsules through 21 days of cold storage at 4□°C

The biological function of the encapsulated product over a 21-day refrigeration period at 4□°C was evaluated by agar-well diffusion assay against *Bacillus cereus* using 0.5 mg/L of the Nisin-chitosan-alginate microcapsule. The results were compared to unrefrigerated controls and unencapsulated Nisin.

#### 2.4.1. Instrumentation, Zeta potential and particle size analysis

Formed microcapsules were diluted in PBS to reduce their viscosity and prevent deterioration before analysis. Dynamic light Scattering (DLS) measurements were obtained using a Zetasizer Pro (Malvern Instruments Ltd., U.K.), which was used to determine the mean particle size, size distribution, intensity average (%) and the polydispersity index (PDI) of the Nisin-alginate-chitosan microcapsules after filtration using a hydrophilic 0.22 μm pore syringe filter. All measurements were taken at 25 °C. Similarly, the surface charge on the microcapsules was determined by measuring the Zeta potential using the Zetasizer Pro. For each sample, the Zetasizer pro equipment was used to automatically take an average of ten (10) readings to obtain a single sample value/reading (Zeta potential), then the mean of three sample readings was utilised to arrive at a final value.

#### 2.4.2. Nisin encapsulation efficiency

Encapsulation efficiency was measured by determining the concentration of free Nisin after encapsulation (unbound Nisin) and comparing this to total Nisin utilised. Following Nisin microcapsule complex formation, the microcapsules were centrifuged at 2000rpm for 30 minutes. The supernatant was carefully decanted from the particles (microcapsules). The supernatant was considered to contain free (unencapsulated) Nisin released during the encapsulation process. The Nisin content of the supernatant was measured first by determining the protein content using the BCA assay and then by HPLC to cross-validate quantification accuracy. For the BCA assay, aliquots of the supernatant were mixed with the BCA working reagent prepared by combining Reagent A and Reagent B in a 50:1 ratio and incubated at 37 °C for 30 minutes. Absorbance was measured at 562 nm using a Jenway spectrophotometer. Protein quantification was performed using a calibration curve generated from bovine serum albumin (BSA) standards, and the concentration of free Nisin was extrapolated from the linear regression equation derived from the standard curve. Encapsulation efficiency was determined using the formula outlined in Equation 3;

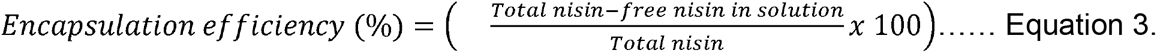

### 2.5. Nisin quantification by HPLC

The concentration of Nisin was quantified using the HPLC method previously described [14] with modifications. A Shimadzu Prominence HPLC equipped with a UV-Photodiode Array Detector was utilised to purify and quantify the Nisin microcapsules. A C18 column (microsob 100-5 Si 250 x 4.6mm) (ThermoFisher Scientific) was used for sample separation. All chemicals used were of analytical grade. The mobile phase employed was a mixture of acetonitrile and methanol acidified with 0.05% Trifluoroacetic acid (TFA) and ultrapure water. Prior to chromatographic run, the microcapsules were precipitated with an equal volume of the mobile phase (matrix matching to avoid denaturation) by centrifugation at 5000rpm for 10 minutes and then filtered using the hydrophilic 0.22 μm syringe filter before injecting into the equipment. This was done to prevent column damage. Chromatographic separation of the sample was undertaken at a column temperature of 40°C with a flow rate of 100µL min^-1^ and an injection volume of 20 µL. The chromatographic method had a total run time of 70 minutes. After an initial hold time of 3 minutes in ultrapure water, a gradient run was undertaken for 60 minutes, wherein the concentration of acetonitrile and methanol was varied from 20% to 75%. This is followed by a 5% column re-equilibration in ultrapure water. UV detection was undertaken at a wavelength of 215nm. Experimental control and result analysis were undertaking using Chromeleon V6.80.

### 2.6. Scanning Electron Microscopy

The morphology of the microcapsules was evaluated by use of a Zeiss EVO15 VP E-Scanning Electron Microscope (Zeiss instruments, Germany). First, the microcapsule was dried using a Labconco Benchtop freeze dryer (Cole Parmer, UK). The dry powdered microcapsules were fixed on to double-sided adhesive carbon sample studs and examined at x4000 magnification and low vacuum at an electron charge of 15kV.

Similarly, SEM was undertaken for treated *Bacillus cereus* to investigate possible cell membrane disruption as the mechanism of action of Nisin 2A. For SEM of *Bacillus cereus,* vegetative cells (before and after treatment with Nisin microcapsule) were fixed in glutaraldehyde to inactivate the organisms. This is followed by dehydration using increasing series of graded ethanol solution, critical point drying with CO_2_ and then gold coated using a gold sputter (thermos Fisher Scientific UK). SEM parameters used for magnification remained the same.

### 2.7. Fourier transform infrared (FTIR) spectroscopy analysis

Complex formation within the microcapsules and the changes in Nisin conformation were evaluated by FT-IR spectroscopy. Spectra of Nisin, sodium alginate, chitosan and Nisin-alginate-chitosan microcapsules were collected. To achieve this, a Bruker IFS 66/s equipped with a single reflection diamond Attenuated Total Reflectance (ATR) accessory (Harrick MVP-Pro™ Star, Bruker Optics Billerica, MA, USA) was utilised at ambient conditions (25 ± 1 °C). Before spectra collection, the sample chamber was flushed with nitrogen gas to limit the impact of water vapour and CO_2_ on the collected spectra. To capture characteristic functional group vibrations of polysaccharides and peptides, FT-IR spectra recording was set to between 400 and 4000cm^-1^ at a resolution of 6cm^-1^, where each spectrum represents an average of 100 scans. For control, background spectra were obtained using the bare diamond crystal while maintaining the same experimental conditions. All obtained spectra were plotted in absorbance units of –log (S/R), where “S” is the single channel spectra of the sample and R that of the bare diamond crystal. Spectral analysis and manipulations were undertaken using the OPUS 6.0 (Bruker Optics Billerica, MA, USA) software.

### 2.8. Assay of Nisin release from the microcapsules *in-vitro*

The Nisin release rate of the microcapsule was determined by suspending 100mg of the microcapsule in 5ml of 0.02M phosphate buffer (pH 5.0), selected to mimic mildly acidic food matrices. This suspension was stirred continuously at 150rpm for 24 hours. At predetermined time points of 2 hours, the supernatant was filtered using a membrane filter. For the determination of eluted Nisin, the protein concentration of the buffer was assayed using the BCA assay protocol and subsequently by reverse-phase HPLC, as described previously. Cumulative release was calculated as a percentage of total encapsulated Nisin.

All experiments were done in triplicates, and results are reported as mean ± standard deviation.

## 3. RESULTS AND DISCUSSION

The physicochemical and biological properties of the synthesized microcapsules were evaluated using scanning electron microscopy (SEM), a Zetasizer for Zeta potential and particle size analysis, Fourier-transform infrared (FTIR) spectroscopy, high-performance liquid chromatography (HPLC). Together, the obtained data provides a comprehensive overview of the formulation’s performance, stability, and potential applications in food preservation.

### 3.1. Impact of calcium concentration on the formation of calcium-alginate pre-gels, particle size and Zeta potential

Relative viscosity measurements were employed to measure pre-gelation of the calcium-alginate pre-gels and the interaction between the calcium divalent cations and the alginate polymer molecules to determine the critical entanglement concentration [15]. The addition of calcium at concentrations ranging from 0 to 2 mM initiated the coiling of the alginate polymer chains, which led to the formation of the microparticle nucleus. Figure 1 presents the effect of calcium concentration on the relative viscosity of calcium-alginate pre-gels.

**Figure 1.**
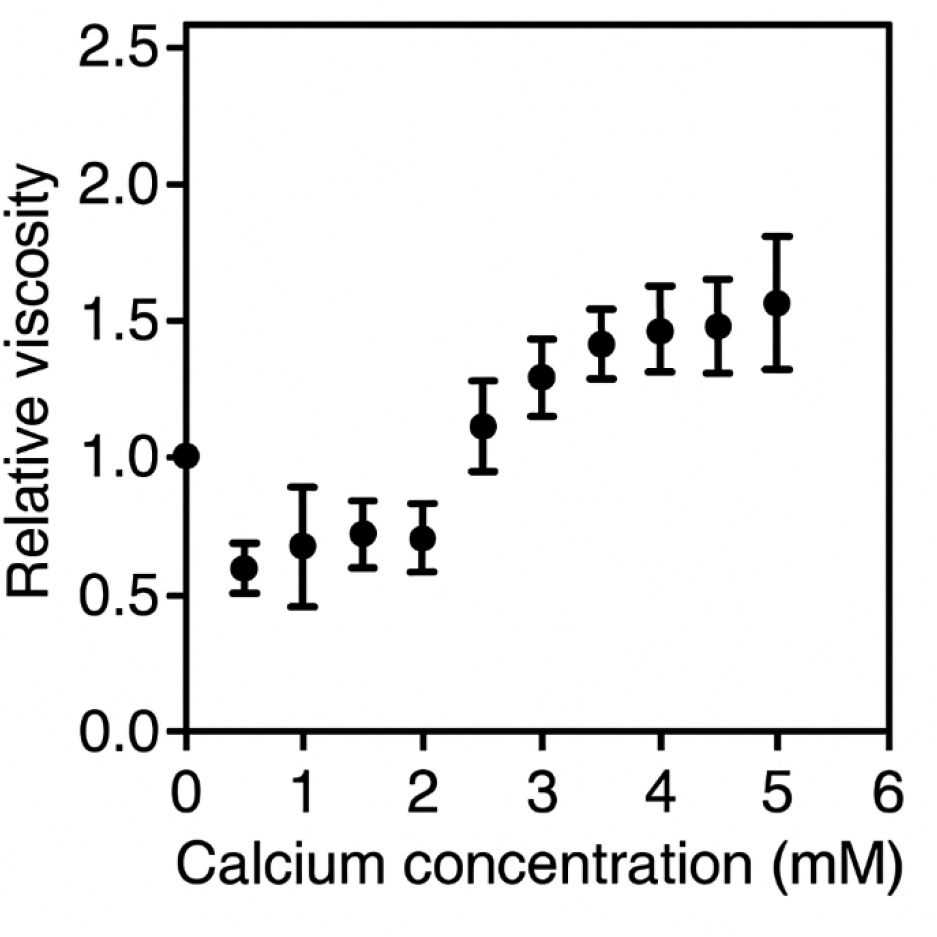
Effect of calcium concentration on the relative viscosity (μr) of calcium-alginate pre-gels

It was observed that at higher calcium concentrations >2mM, there is an increase in the relative viscosity of the pre-gels (Figure 1), possibly due to intermolecular interactions between alginate chains, facilitated by the binding of calcium ions where they act as cross-linking agents, helping to form ionic bridges between the residues of guluronic acid in the alginate chains, eventually leading to a full-gelled state [16]. This has been demonstrated in a previous study by Li et al. [17] which showed that calcium ions significantly influences the gelation kinetics and mechanical properties of alginate-based gels by enhancing the network structure of polymer through crosslinking. Conversely, a reduction in the calcium concentration to 2mM brings about a reduction in the relative viscosity of the sodium alginate solution and this reduced viscosity through a lower Ca^2+^ concentration has been shown to bring about intramolecular crosslinking of individual polymer chains independent of intermolecular linkages, leading to the formation of compact coiled chains and reduced viscosity [18], which is ideal for the formation of microcapsules. The findings in this present study correlates with previous observations that the size and Zeta potential of microparticles is optimised with lower calcium concentrations which enhances the formation of smaller sized particles and more uniform Zeta potential values, thereby reducing aggregation and ensuring that the encapsulation process is better controlled, as exemplified in a previous study by Chandrasekar et al. [8] which found that calcium concentration had a significant effect (p < 0.05) on particle size. Similarly, a more recent study by Xu et al. [19] demonstrated that the intramolecular cross-linking reaction of alginate with Ca^2+^ produced a compact coiled structure and pre-gel state at low ionic concentrations, resulting in reduced particle sizes, whereas too high a concentration of Ca^2+^ may cause cross-linking of alginate chains laterally, resulting in bigger structures and aggregation.

### 3.2. Impact of pH and pKa values on Microcapsule formation

Following pre-gelation, Nisin-alginate-chitosan microcapsules were synthesised by adding Nisin solution dropwise into the sodium alginate pre-gel and then chitosan solution. As has been previously outlined by literature [20], microcapsules formation is affected mainly by the pH of formation, calcium and chitosan concentration. In this present study, optimal pH of encapsulation is 5.0. This corresponds with the findings of a recent work investigating alginate–chitosan complexes formed with different polymers [21]. In this study, it was demonstrated that pH of formation influences the interaction between chitosan and alginate as these interactions are governed by the dissociation constants (pKa values) of the polymers. Maintaining pH between the dissociation constants of chitosan and alginate (6.5-6.6 for chitosan, 3.38 and 3.65 for the pKa values of mannuronic and guluronic acid monomers in alginate respectively), the anionic carboxyl groups of alginate and the cationic amino groups of chitosan are ionized, and polyelectrolyte complexation is optimized, producing microcapsules with higher payload retention and narrower size distributions. Similarly, another study by Vahedifar et al. [22] further supports this by highlighting that calcium ions facilitate ionic crosslinking within the alginate pre-gel at optimal pH, while the concentration of chitosan determines the extent of polymer interaction, which collectively influence the stability and compactness of the microcapsules as observed in this present study.

The protonation state of chitosan’s amine groups and the deprotonation of alginate’s carboxyl groups are pH-dependent, influencing their ability to interact and form a stable matrix. At suboptimal pH or at high chitosan concentrations, aggregation and poor encapsulation efficiency were observed, which supports earlier studies indicating that fine-tuning of pH, calcium ion concentration, and polymer ratios is critical for successful encapsulation and the overall structural stability of the encapsulated bioactive agents, in this case, Nisin. For example, Bustos et al. demonstrated that the structural stability of calcium–alginate complexes is strongly dependent on both pH and calcium ion concentration, with lower Ca²⁺ levels and acidic conditions promoting more stable polymeric networks and slower release of encapsulated bioactive compounds, whereas deviations from these conditions led to reduced structural integrity and enhanced release [23]. In this present study, pH was carefully modulated across the different experimental runs to ensure optimal ionization of functional groups in both chitosan and alginate, which is critical for the ionic gelation and polyelectrolyte complexation mechanisms central to microcapsule formation. Chitosan, a polycationic biopolymer, has a pKa of ∼6.5, and its amine groups must be protonated (–NH□□) for electrostatic interaction with the carboxylate groups (–COO□) of alginate, which are deprotonated above pH ∼3.4–3.6 (the pKa range for guluronic and mannuronic acid residues). Thus, a moderately acidic pH (typically between 4.5 and 5.5) is optimal to achieve sufficient ionization of both polymers and enable stable complex formation. In this study, encapsulation was conducted at such a pH (5.0), resulting in microcapsules with good structural integrity. In the current study, deviations from optimal pH conditions led to noticeable changes in microcapsule performance. When chitosan concentration was increased to 1%, a corresponding shift in pH was observed, resulting in a PDI of 1.0 and a significant drop in encapsulation efficiency, as well as broad size distribution. This was likely due to over-protonation of chitosan or suboptimal ionization of alginate, leading to excessive intra- and intermolecular interactions and reduced availability of functional sites for Nisin entrapment. This is supported by the seminal work of Shah et al. which demonstrated that both pH and chitosan concentration play a critical role in governing molecular interactions during microcapsule synthesis, with excess cationic charge promoting particle aggregation and reducing peptide encapsulation efficiency [24]. It was found that higher pH of 5.0-7.0 resulted in weak electrostatic repulsion between droplets and the formation of larger and possibly aggregated microcapsules. Thus, maintaining pH in the optimal range <5.0 during chitosan-based nanoparticle formation was critical for encapsulation efficiency and stability under food-relevant conditions. These results showed that minor deviations in pH (from 4.0 to 5.0-7.0) led to a substantial decrease in encapsulation efficiency, supporting the conclusion that the protonation state of chitosan’s amine groups is a limiting factor in its interaction with other polyanions and payload molecules. This is particularly relevant in food systems where ionic strength and pH vary significantly such as in fruit juice matrixes.

### 3.3. Zeta potential of Nisin-loaded microcapsules measurements

Dynamic Light Scattering (DLS) measurements of the microcapsules revealed that they have an average size of 150nm (Figure 2) which agree with the findings of previous studies such as Lee et al. [4], in which chitosan and N-Cs nanoparticle mean size of 64.34 and 147.93nm was obtained respectively. Similarly, the obtained microcapsules were polydispersed with a polydispersity index (PDI) of 0.302. A PDI of 0.302 point to a relatively uniform size distribution which is important for slow-release profiles and consistent antimicrobial performance. Furthermore, this PDI suggests a relatively narrow size distribution, reflecting consistent particle formation and good formulation control, similar to the 0.51 obtained by Pinilla et al. [25]. Overall, this points to a well-dispersed system with strong electrostatic stabilization which is ideal for applications in food systems due to increased surface area-to-volume ratios, improved bioavailability, and efficient interaction with microbial membranes [25].

**Figure 2.**
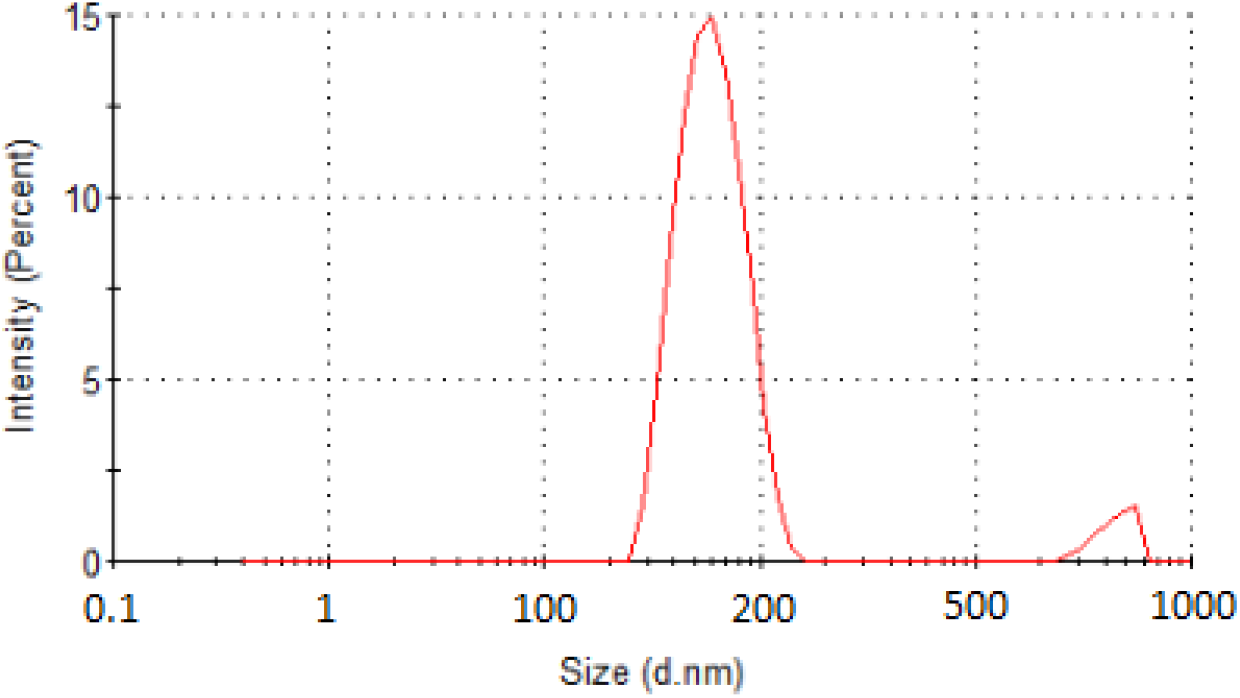
Size distribution of Nisin-loaded microcapsule by Dynamic Light Scattering (DLS)

Also, the microcapsules have a Zeta potential of +36.4mV. This positive Zeta potential indicates that the surface of the microcapsule is positively charged, ostensibly due to the presence of positively charged amino acid groups on the surface [26]. The Zeta potential impacts the microcapsule’s stability due to the electrostatic interaction and repulsion between the particles. It indicates the charge of the microcapsules and can serve as an indicator of their stability. Zeta potential values above ±30 mV are typically considered stable, reducing the risk of aggregation during storage, as higher absolute Zeta potential values generate electrostatic repulsion strong enough to prevent particle aggregation [27]. The obtained Zeta potential of +36.4mV in this present study indicates a strong surface charge repulsion, which helps prevent aggregation of microcapsules and enhances colloidal stability by promoting electrostatic repulsion between particles. This result is comparable to the Zeta potential of +39.4mV reported previously by Khan et al. [28] and +33.4mV in a related study by Lee et al. [4] for similar chitosan-based encapsulated Nisin systems. These comparable surface charge values further confirm that the encapsulation approach used in the present study leads to a stable formulation suitable for suspension in aqueous food matrices like juice or milk.

Of note is that concentration of chitosan significantly affected the size and polydispersity of the microcapsule, with multiple peaks recorded in the DLS analysis of the chitosan-Nisin-alginate microcapsules when the concentration of chitosan in the formulation was increased to 1% and a ratio of 1:1. Furthermore, as highlighted previously, increasing the chitosan concentration resulted in monodispersed microcapsule with a PDI of 1.0 thus indicating likely aggregation. This is attributable to the lower encapsulation efficiency of this combination, thus, chitosan of 0.5% was utilised to improve the polydispersity, as encapsulation efficiency and polydispersity are highly dependent on the solution’s pH and chitosan concentration [20]. The observed changes in PDI based on chitosan concentration highlights the importance of carefully balancing polymer concentration and ionic crosslinking for optimal encapsulation outcomes and mirrors the observations made in earlier studies by Jin et al. [29] and Isibor [30], where it was found that increasing chitosan content beyond optimal thresholds resulted in broader size distributions and decreased encapsulation efficiency due to excessive polymer–polymer interaction and reduced crosslinking homogeneity. This is important as the physicochemical characteristics of the microcapsules including particle size distribution, colloidal stability, and surface charge directly influence functionality, dispersion in food matrices, and storage behaviour. Similarly, the lower encapsulation efficiency observed at higher chitosan concentrations may be attributed to the increase in intermolecular interactions within the chitosan-Nisin-alginate microcapsules, which possibly hindered effective Nisin incorporation, coupled with the crucial role of the pH solution in the protonation of the chitosan amino group and its effect in the interaction with the encapsulated compound. The study by Shah et al. [24] demonstrated that optimizing the chitosan concentration and maintaining a favourable pH and storage time, significantly improved encapsulation efficiency and resulted in uniform microcapsules with improved stability against pH and salts, and functional performance in the delivery of bioactive compounds [24]. Hence, this present study’s use of higher calcium concentrations (2mM) for modifications of the chitosan-Nisin microcapsules as calcium ions have been demonstrated to facilitate crosslinking, influence gelation kinetics and stabilize microcapsule structure [8, 31].

### 3.4. Overview of encapsulation modelling following the Response Surface Methodology

The effects of chitosan (CS) concentration, relative viscosity of the calcium-alginate-Nisin solution, and CS/Nisin-alginate ratio on Zeta potential were examined through a second-order regression model. Zeta potential, a critical factor for colloidal stability and electrostatic interactions, varied between 34.2 mV and 38.9 mV across the 15 experimental runs (Table 2). Each run represented a unique combination of these formulation variables, enabling the model to explore the full design space while minimizing the number of experiments. This design allowed for the detection of both linear and interaction effects between the variables on encapsulation efficiency, Zeta potential, and PDI. Among the fifteen runs, Run 10, characterized by a chitosan concentration of 1.0 mg/mL, relative viscosity of 1.0, and a CS:Nisin–alginate ratio of 1:2, produced the highest encapsulation rate (65.0%) with a favourable Zeta potential (36.4 mV) and the lowest polydispersity index (0.302).

**Table 2.** Experimental values of response variables for response surface design.

| Experimental Run | Variables |  |  | Encapsulation Rate (%) | Zeta potential (mV) | PDI |
| --- | --- | --- | --- | --- | --- | --- |
|  | CS Concentration (mg/mL) | Relative Viscosity | CS/Nisin-alginate Ratio |  |  |  |
| 1 | 0.5 | 0.8 | 1:1 | 58.3 | 34.2 | 0.445 |
| 2 | 0.5 | 1.2 | 1:1 | 60.1 | 35.5 | 0.402 |
| 3 | 0.5 | 1.0 | 1:1 | 61.0 | 36.0 | 0.390 |
| 4 | 1.0 | 0.8 | 1:2 | 63.5 | 37.5 | 0.365 |
| 5 | 1.0 | 1.2 | 1:2 | 64.2 | 38.2 | 0.325 |
| 6 | 1.0 | 1.0 | 1:2 | 63.0 | 37.0 | 0.341 |
| 7 | 1.5 | 0.8 | 2:1 | 59.5 | 34.9 | 0.410 |
| 8 | 1.5 | 1.2 | 2:1 | 60.7 | 35.7 | 0.388 |
| 9 | 1.5 | 1.0 | 2:1 | 61.8 | 36.5 | 0.375 |
| 10 | 1.0 | 1.0 | 1:2 | 65.0 | 36.4 | 0.302 |
| 11 | 0.75 | 1.0 | 1:2 | 62.5 | 36.8 | 0.340 |
| 12 | 1.25 | 1.0 | 1:2 | 62.9 | 37.2 | 0.348 |
| 13 | 1.0 | 0.9 | 1:2 | 64.3 | 38.5 | 0.318 |
| 14 | 1.0 | 1.1 | 1:2 | 64.7 | 38.9 | 0.310 |
| 15 | 1.0 | 1.0 | 1:1 | 62.0 | 36.2 | 0.360 |

The regression coefficient and summary (Table 3, 4 and 5) revealed that both the linear and quadratic terms for CS concentration had near-threshold p-values (p = 0.0748 and p = 0.0757, respectively), suggesting a possible non-linear influence of CS dosage on particle surface charge. While these results were not statistically significant at the conventional 5% level, they indicate a moderate trend that may warrant further investigation.

**Table 3.** Regression Coefficients for Encapsulation Rate Model.

| Term | Estimate | Std. Error | t-value | p-value |
| --- | --- | --- | --- | --- |
| (Intercept) | 1.804 | 22.358 | 0.081 | 0.9388 |
| CS Concentration (Linear) | 77.241 | 30.308 | 2.549 | 0.0514 |
| CS Concentration (Quadratic) | -30.046 | 11.651 | -2.579 | 0.0495 |
| Relative Viscosity (Linear) | 59.620 | 27.634 | 2.157 | 0.0835 |
| Relative Viscosity (Quadratic) | -27.672 | 13.627 | -2.031 | 0.0980 |
| CS/Nisin Ratio (Linear) | -12.328 | 5.667 | -2.175 | 0.0816 |
| CS/Nisin Ratio (Quadratic) | 20.680 | 9.318 | 2.219 | 0.0772 |
| CS Conc × Rel. Viscosity (Interaction) | -3.450 | 5.442 | -0.634 | 0.5540 |
| CS Conc × CS/Nisin Ratio (Interaction) | -25.798 | 13.664 | -1.888 | 0.1177 |
| Rel. Viscosity × CS/Nisin Ratio (Interaction) | 1.950 | 3.416 | 0.571 | 0.5928 |

**Table 4.** Regression Coefficients for Zeta potential Model.

| Term | Estimate | Std. Error | t-value | p-value |
| --- | --- | --- | --- | --- |
| (Intercept) | -14.120 | 22.248 | -0.635 | 0.5536 |
| CS Concentration (Linear) | 67.693 | 30.158 | 2.245 | 0.0748 |
| CS Concentration (Quadratic) | -25.908 | 11.594 | -2.235 | 0.0757 |
| Relative Viscosity (Linear) | 47.773 | 27.498 | 1.737 | 0.1428 |
| Relative Viscosity (Quadratic) | -22.155 | 13.560 | -1.634 | 0.1632 |
| CS/Nisin Ratio (Linear) | -10.169 | 5.639 | -1.803 | 0.1312 |
| CS/Nisin Ratio (Quadratic) | 20.340 | 9.272 | 2.194 | 0.0797 |
| CS Conc × Rel. Viscosity (Interaction) | -2.075 | 5.415 | -0.383 | 0.7173 |
| CS Conc × CS/Nisin Ratio (Interaction) | -26.005 | 13.597 | -1.913 | 0.1140 |
| Rel. Viscosity × CS/Nisin Ratio (Interaction) | 0.825 | 3.399 | 0.243 | 0.8179 |

**Table 5.**
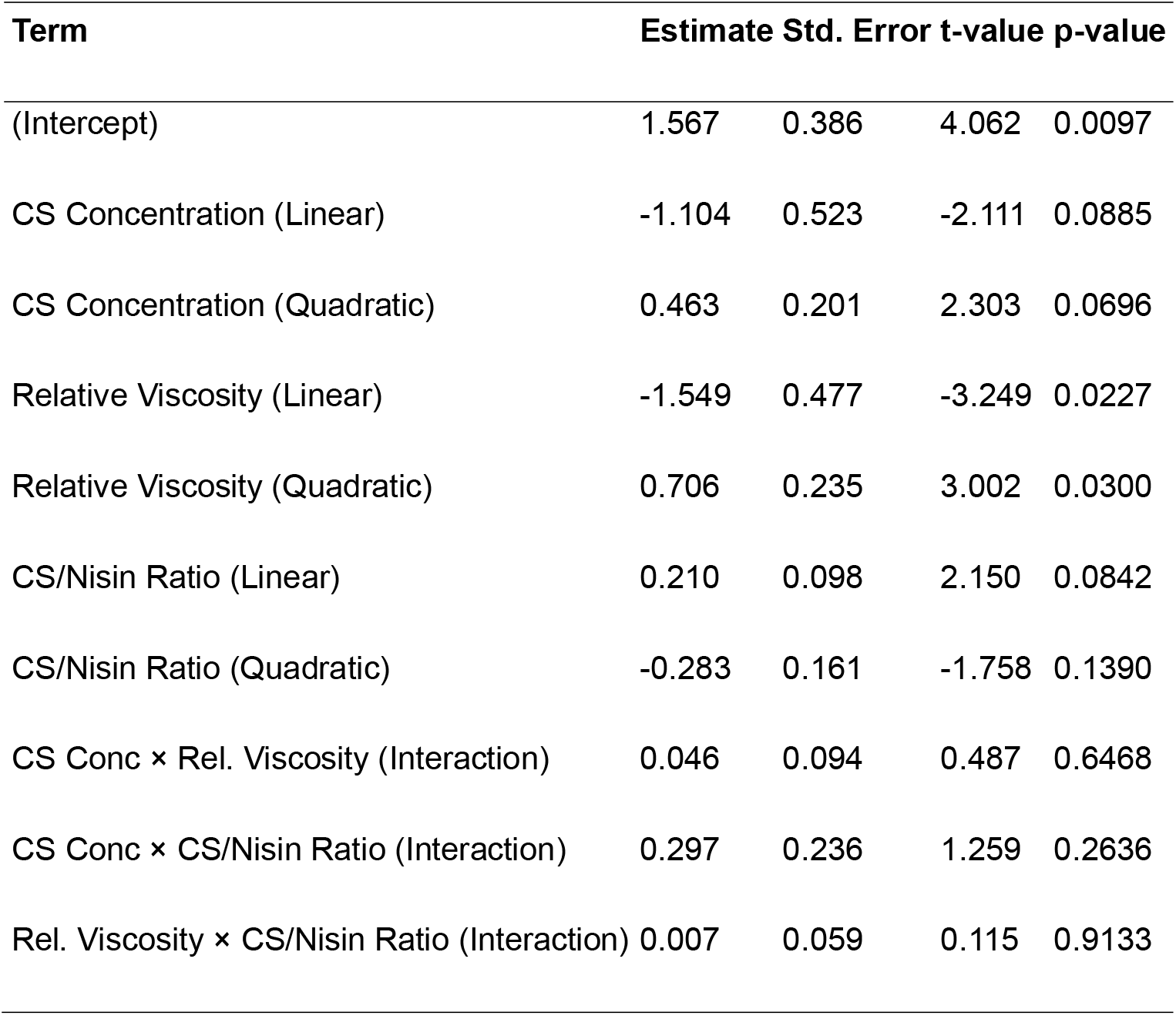
Regression Coefficients for PDI Model.

| Term | Estimate | Std. Error | t-value | p-value |
| --- | --- | --- | --- | --- |
| (Intercept) | 1.567 | 0.386 | 4.062 | 0.0097 |
| CS Concentration (Linear) | -1.104 | 0.523 | -2.111 | 0.0885 |
| CS Concentration (Quadratic) | 0.463 | 0.201 | 2.303 | 0.0696 |
| Relative Viscosity (Linear) | -1.549 | 0.477 | -3.249 | 0.0227 |
| Relative Viscosity (Quadratic) | 0.706 | 0.235 | 3.002 | 0.0300 |
| CS/Nisin Ratio (Linear) | 0.210 | 0.098 | 2.150 | 0.0842 |
| CS/Nisin Ratio (Quadratic) | -0.283 | 0.161 | -1.758 | 0.1390 |
| CS Conc × Rel. Viscosity (Interaction) | 0.046 | 0.094 | 0.487 | 0.6468 |
| CS Conc × CS/Nisin Ratio (Interaction) | 0.297 | 0.236 | 1.259 | 0.2636 |
| Rel. Viscosity × CS/Nisin Ratio (Interaction) | 0.007 | 0.059 | 0.115 | 0.9133 |

The model equation for Zeta potential is expressed as:

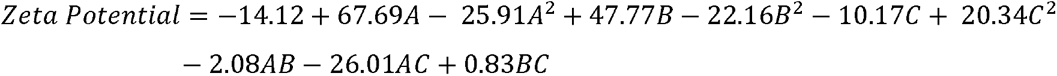

Where A, B, and C are the coded variables representing CS concentration, relative viscosity, and CS/Nisin-alginate ratio, respectively.

The interaction terms were not statistically significant, suggesting that the factors acted independently on Zeta potential within the studied ranges. The 3D response surface (Figure 3) shows a smooth convex region, with the maximum Zeta potential observed at CS = 1.0 mg/mL, viscosity = 1.0, and CS: Nisin = 1:2, which coincides with Run 10, suggesting that it is the optimal formulation for surface charge stability.

**Figure 3.**
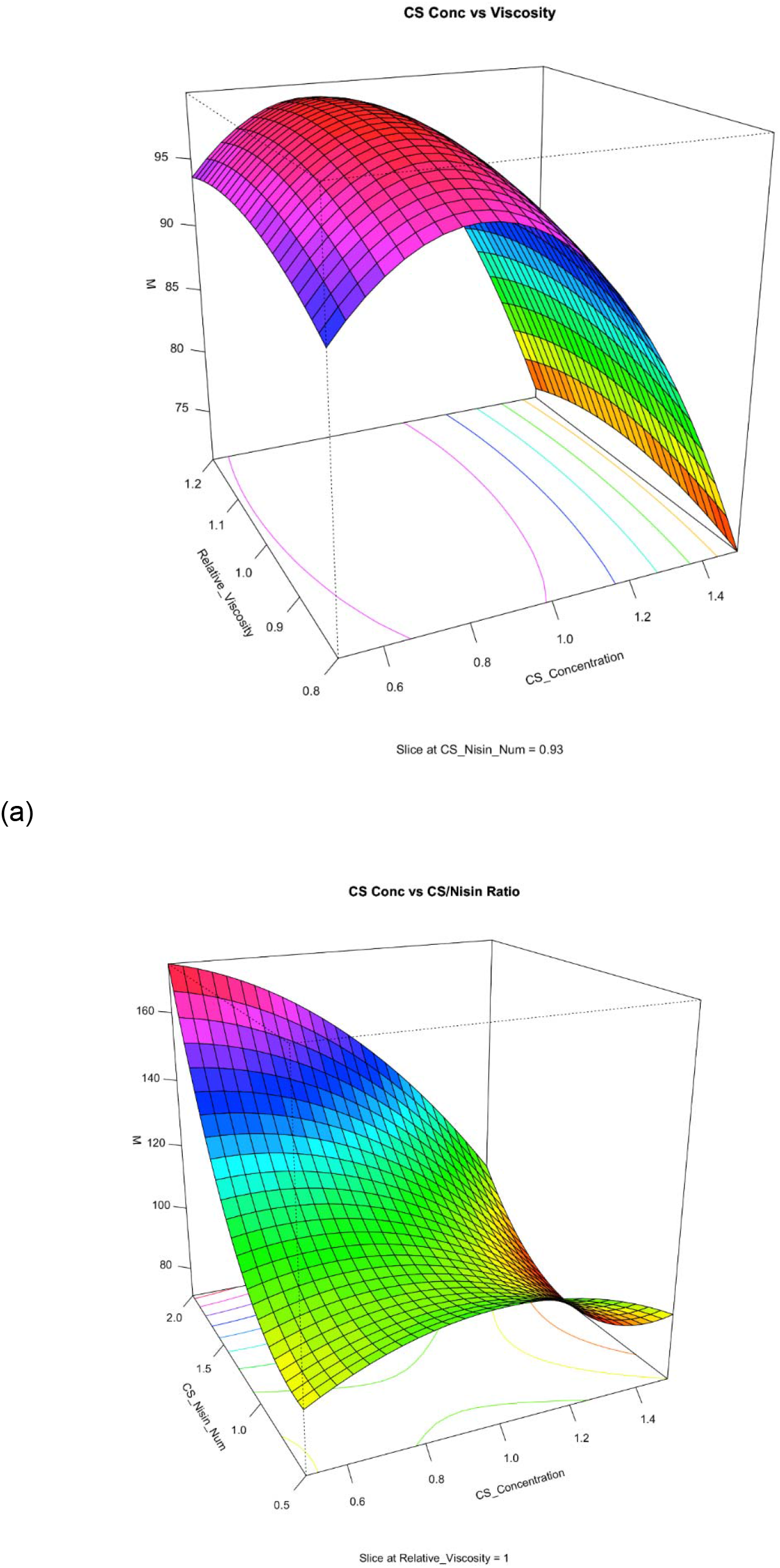
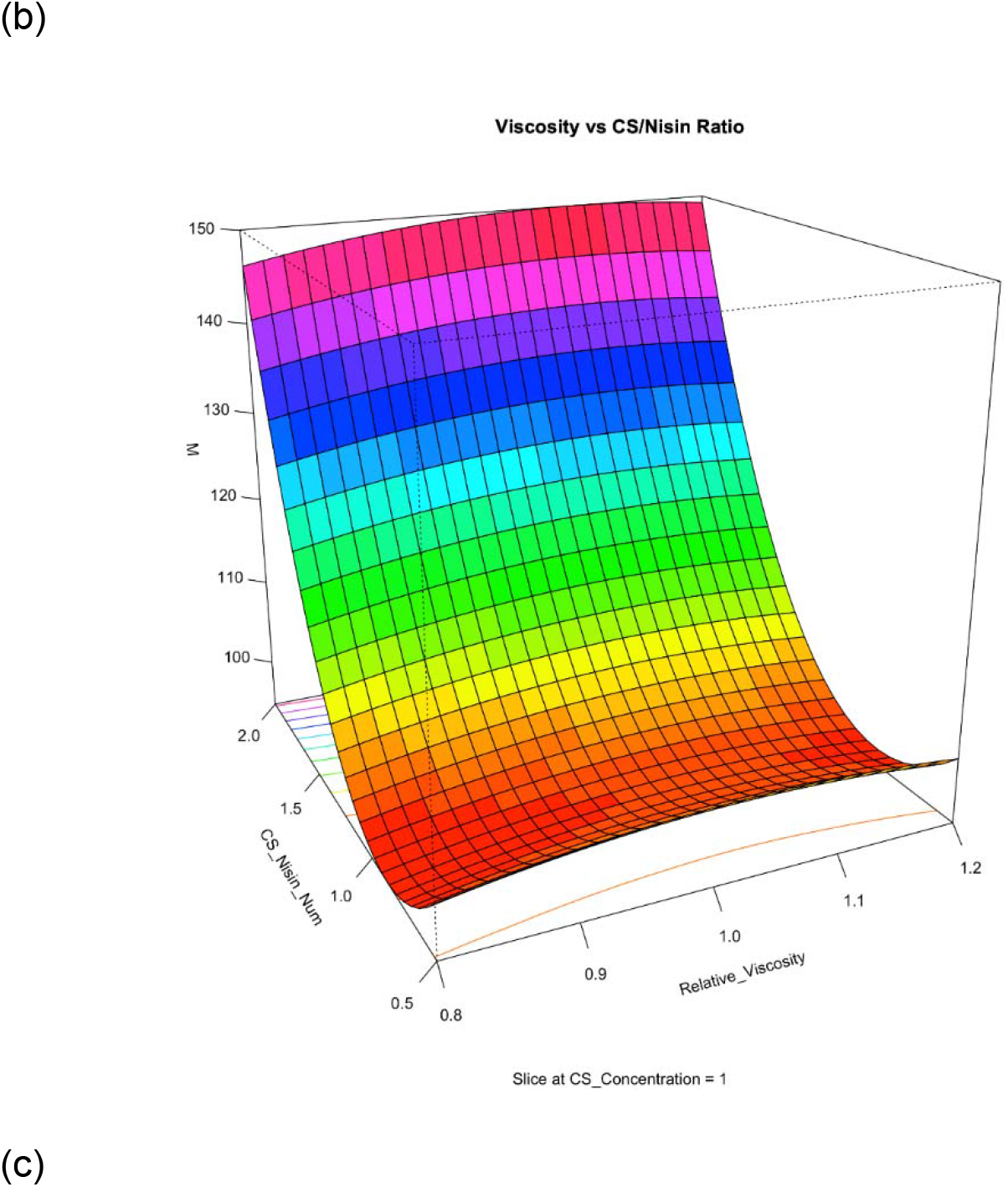
Response surface plots showing the interaction effects of formulation parameters on the composite response (M = encapsulation rate + Zeta potential). (a) CS concentration vs Relative Viscosity (b) CS concentration vs CS/Nisin ratio (c) Relative Viscosity vs CS/Nisin ratio

In contrast, PDI values ranged from 0.302 to 0.445, reflecting a narrow and acceptable size distribution. From the regression summary (Table 3, 4 and 5), relative viscosity had a significant linear (*p* = 0.0227) and quadratic (*p* = 0.0300) influence on PDI, while CS concentration and CS/Nisin ratio showed weaker effects. Despite some near-significant trends, interaction terms did not significantly affect PDI. These findings suggest that all formulations produced relatively homogenous microcapsules, with optimal uniformity at Run 10 (PDI = 0.302).

### 3.5. Encapsulation Efficiency Modelling

Encapsulation efficiency, here referred to as the encapsulation rate, is crucial for quantifying the loading capacity of Nisin within the chitosan-alginate network. The observed encapsulation rates ranged from 58.3% to 65.0%, with the highest rate recorded at Run 10 under conditions of CS = 1.0 mg/mL, viscosity = 1.0, and CS/Nisin = 1:2 (shown earlier in Table 2).

The regression model for encapsulation efficiency was statistically robust (*R²* = 0.9361, *p* < 0.05), and the fitted equation is given as:

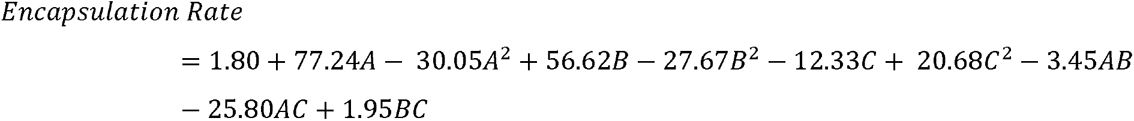

Significant predictors included the quadratic term of CS concentration (*p* = 0.0495), with near-significant effects from the linear terms of CS and viscosity (*p* = 0.0514 and 0.0835, respectively) (Table 3, 4 and 5). These results suggest a peak encapsulation performance at intermediate levels of CS and viscosity.

### 3.6. Optimisation of the Chitosan-Nisin-Alginate System

To achieve a balanced formulation with both high encapsulation and desirable electrostatic stability, a composite response variable (M) was defined as:

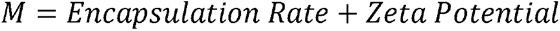

Regression modelling of M yielded the following fitted equation

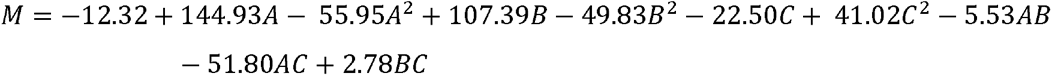

The composite model had a high multiple R-squared of 0.9184, indicating good explanatory power. Although not all terms were statistically significant at *p* < 0.05, the key effects of CS concentration and CS/Nisin ratio were near significant (Table 6), and their inclusion was justified by improved model fit and interpretability.

**Table 6.** Regression Coefficients for Composite Metric (Encapsulation + Zeta potential)

| Term | Estimate | Std. Error | t-value | p-value |
| --- | --- | --- | --- | --- |
| (Intercept) | -12.316 | 43.876 | -0.281 | 0.7902 |
| CS Concentration (Linear) | 144.934 | 59.477 | 2.437 | 0.0589 |
| CS Concentration (Quadratic) | -55.954 | 22.864 | -2.447 | 0.0581 |
| Relative Viscosity (Linear) | 107.393 | 54.229 | 1.980 | 0.1045 |
| Relative Viscosity (Quadratic) | -49.828 | 26.742 | -1.863 | 0.1215 |
| CS/Nisin Ratio (Linear) | -22.497 | 11.122 | -2.023 | 0.0990 |
| CS/Nisin Ratio (Quadratic) | 41.020 | 18.287 | 2.243 | 0.0749 |
| CS Conc × Rel. Viscosity (Interaction) | -5.525 | 10.680 | -0.517 | 0.6270 |
| CS Conc × CS/Nisin Ratio (Interaction) | -51.802 | 26.815 | -1.932 | 0.1112 |
| Rel. Viscosity × CS/Nisin Ratio (Interaction) | 2.775 | 6.703 | 0.414 | 0.6960 |

The response surface plots provided visual insight into the interaction dynamics. Figure 3a (CS Conc vs Viscosity) revealed a convex surface with a maximum near x1 = x2 = 0, corresponding to CS = 1.0 and viscosity = 1.0 (A well-defined saddle-shaped surface with an optimal region near the center (coded levels ≈ 0), indicating that encapsulation rate increases with CS and viscosity up to a point, after which it declines, likely due to polymer saturation and diffusion limitations). Figure 3b (CS Conc vs CS:Nisin Ratio) and Figure 3c (Viscosity vs CS:Nisin Ratio) similarly highlighted a peak around CS:Nisin = 1:2 (x3 = -1).

The experimental and predicted optimum was identified at:

- CS concentration = 1.0 mg/mL
- Relative viscosity = 1.0
- CS:Nisin ratio = 1:2

This condition corresponded to:

- Encapsulation rate = 65.0%
- Zeta potential = 36.4 mV
- Composite score M = 104.4

The result revealed a unique trend: the encapsulation rate increased with increasing CS concentration and solution viscosity but only to a limit of 1mg/ml. At more than moderate concentrations (1.5mg/ml), the encapsulation efficiency started to diminish. The trend is in line with previous reports that show that very high concentrations of polymer could hinder the even distribution of active agents due to increased solution viscosity and entanglement of polymers [32]. In this case, the highest encapsulation rate of 65% was reported at the concentration of 1.0 mg/mL CS and the relative viscosity of 1.0, i.e., moderate concentration favours symbiotic interaction between biopolymer and active ingredient. Interestingly, the CS: Nisin-alginate ratio also significantly contributed, though in a less striking way. The 1:2 ratio produced the optimum result for encapsulation as well as Zeta potential, suggesting that more Nisin-alginate than chitosan allowed greater encapsulation while preserving good electrostatic characters. This could be due to greater ionic compatibility between Nisin and the alginate matrix with reduced domination by the chitosan chains.

The Zeta potential values ranged from 34.2 mV to 38.9 mV, and the highest value was at the 1:2 ratio again. These results are unique and significant since surface charge influences the shelf life and stability of colloidal systems. A higher Zeta potential tends to indicate a higher repulsive force between particles, and this inhibits aggregation in the long run [27]. Even if statistical significance was weak in all variables in the Zeta potential model, experimental trends agreed with each other and were of biological relevance. As for polydispersity, the index remained below 0.45 for all samples, which reflects a relatively narrow particle size distribution. This observation supports the idea that the formulations, particularly those near the optimum, were homogenous and well-formed. Notably, viscosity emerged as the most influential factor on PDI, with higher values leading to slightly broader distributions. This aligns with the common challenge of increased heterogeneity at higher viscosities due to less efficient mixing during the encapsulation process.

By combining encapsulation rate and Zeta potential into a single optimisation metric (M), it was possible to visualise how all three formulation variables interact simultaneously. The 3D response surfaces showed that the optimal condition, 1.0 mg/mL CS, 1.0 relative viscosity, and a 1:2 CS: Nisin-alginate ratio, sat at the peak of all model predictions. The experimental results obtained (65% encapsulation efficiency) agrees with the predictions of the model for peak encapsulation performance at intermediate levels of CS and viscosity, which adds credibility to the modelling approach used. These findings suggest that the system is not only statistically robust but also practically reliable.

Most of the variation was predicted by the models, yet some p-values remained greater than the common threshold (0.05). However, the overall predictive patterns were stable and aligned well with experimental observations. This indicates that models are here to support understanding and not replace consistent findings in research. While a low p-value can suggest potentially significant effects, the true strength of this work lies in the consistency seen in all the results. Encapsulations and Zeta potentials predicted and measured closely tracked one another, regression models exhibited similar trends, and response surface plots all pointed to the same best formula. This overall consistency gives a more confident outcome than any single individual statistical test in isolation.

### 3.7. Encapsulation efficiency and yield of Nisin-chitosan-alginate microparticles

For each batch of microparticles prepared using 125 ml of 0.05% chitosan, 250 ml of 0.05% alginate, 25 ml of 2 mM calcium chloride and 400 mg Nisin, a theoretically assumed mass of 600mg per batch was assumed, provided that all the added polymers were utilised in the formation of the microparticles. However, the microparticles’ actual yield and encapsulation efficiency was determined using Equation (3) as previously highlighted and was determined to be 65%, indicating that some of the material utilised for the encapsulation at the formation stage wasn’t utilised in microcapsule formation and was lost to the supernatant. This can be attributed to the ratios of chitosan to alginate, which is important as interactions between chitosan and alginate polymers are dependent on the molecular weight of the polymers [8] and the concentration of calcium within the alginate pre-gel [33]. Overall, the polyelectric films containing a ratio of 1:2 of chitosan to alginate (without calcium) had the highest level of complexation between the negatively charged carboxyl groups in alginate and positively charged amine groups in chitosan [8]. At higher calcium concentrations, there was increased binding of some of the negatively charged side chains between chitosan and alginate, resulting in fewer interaction and reduced encapsulation efficiency. This finding corroborates the results obtained in a recent study by Xu et al. [34] where increasing calcium concentration beyond 7.5 mM, impacted the stability of the hydrogel and reduced the thermodynamic stability of the ionic crosslinks, which resulted in low encapsulation of lutein used in their study. Overall, the total amount of Nisin utilised for batch production of microparticles was 10 mg for every 80 mg/g of alginate. Following encapsulation, it was determined that the concentration of Nisin in the supernatant after centrifugation using the Pierce BCA assay was 3.50 ± 0.5 mg, thus establishing an encapsulation efficiency of 65%.

This value is comparable to microparticle encapsulation efficiency values of Nisin reported by Hosseini et al. [35] in which the encapsulation efficiency of Nisin microcapsules encased in alginate-starch was between 48.33 and 54.58%, and this is dependent on the ratio of Nisin/alginate utilised (%w/w) [35]. Similarly, as reported by Bernela et al. [11], an encapsulation efficiency of 41.45 to 88.36% was obtained for Nisin in alginate–chitosan–pluronic composite nanoparticles under various treatment conditions of peptide to polymer ratio, cross-linker concentration, Nisin concentration and stirring speed [11].

Furthermore, one of the most important considerations in designing such delivery systems is the delicate balance between encapsulation efficiency and microcapsule stability. Improved encapsulation efficiency yields higher payload of bioactive material in the carrier matrix, while excessive loading levels or dense polymer cross-linking may compromise the structural uniformity, leading to premature leakage or brittle particles [36]. Conversely, while increased microcapsule rigidity which could be caused by tighter ionic cross-linking or higher concentrations of chitosan can enhance the physical stability and stress resistance of the microcapsule, it can degrade controlled release kinetics and diminish bioavailability of the entrapped peptide [37]. The optimized formulation in this study (65% encapsulation efficiency; +39.4 mV zeta potential) shows an ideal balance between these parameters, being both highly stable and highly efficient in entrapment. Similar studies on encapsulation of bacteriocin or protein in chitosan–alginate systems have shown that being electrostatically balanced and possessing low ratios of the polymers ensures microcapsule integrity without compromising diffusion-facilitated release [38, 39]. Thus, the efficacy of this system relies on the achievement of a balance of polymers that will ensure maximum encapsulation with flexibility as well as long-term functionality during storage and use in acidic drinks.

### 3.8. HPLC analysis of Encapsulated Nisin 2A following release

High-performance liquid chromatography (HPLC) remains the gold standard for the qualitative and quantitative assessment of antimicrobial peptides such as Nisin, owing to its high sensitivity, precision, and ability to detect subtle structural changes [40, 41]. In this study, HPLC analysis was employed to evaluate the structural integrity of Nisin following its release from the chitosan-alginate microcapsules. The retention times of Nisin following encapsulation were recorded at 28.148 and 28.752 minutes (Figure 4), indicating the presence of distinct yet closely eluting peaks, which are consistent with the behaviour of pure Nisin. This result is similar to the HPLC results reported by a previous study on Nisin by Slootweg et al. [42] in which two peaks were obtained with a retention time of 25.97 and 26.17, respectively, with a combined peak area of 1528.7. Figure 4 shows the chromatogram of Nisin with a retention time of 28.148 and 28.752 minutes.

**Figure 4.**
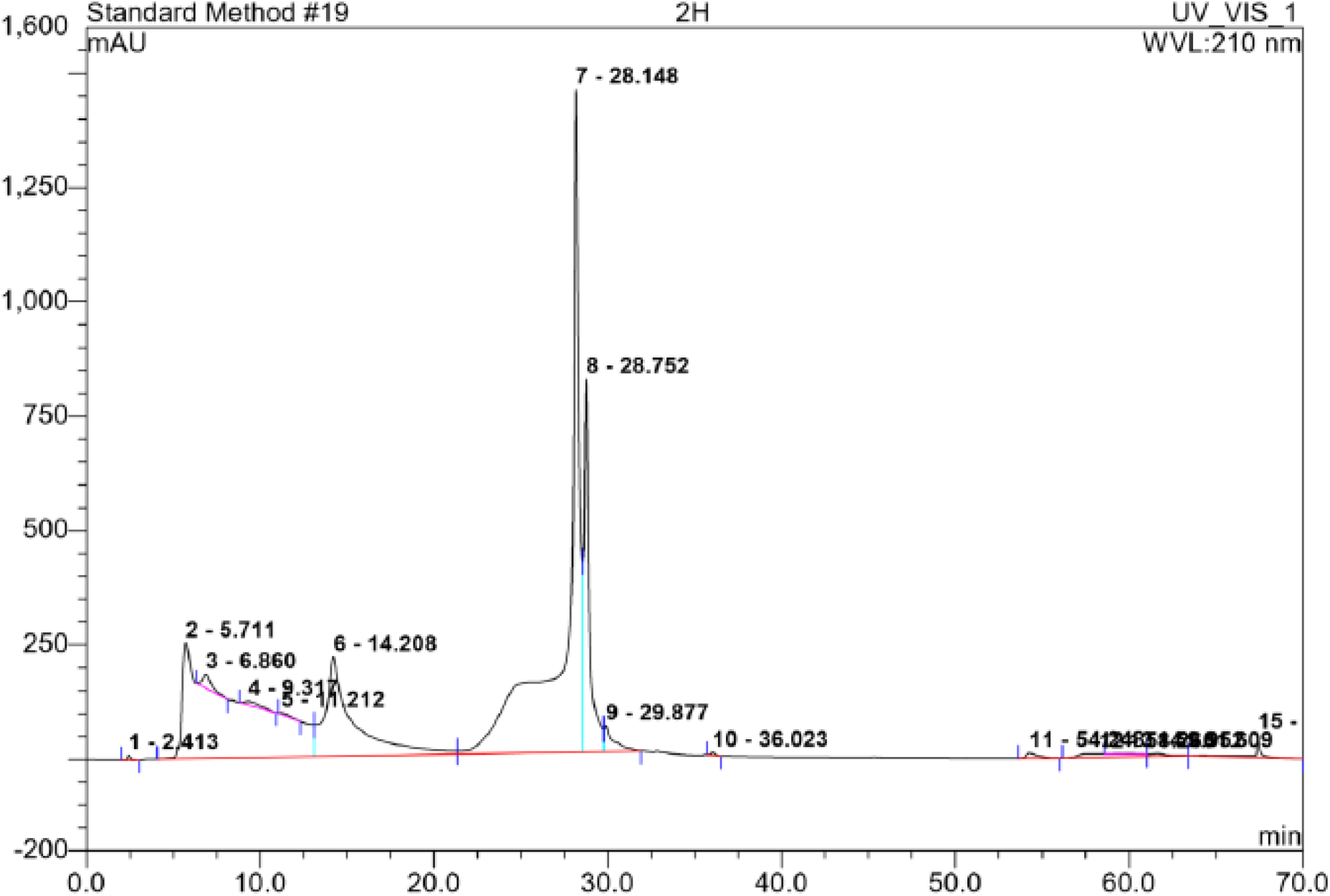
HPLC chromatograms of Nisin 2a following encapsulation

Although there is a slight variation in retention time values between the present study and earlier studies [42], this discrepancy can be attributed to differences in column type, mobile phase composition, gradient program, and column temperature. In this present study, a C18 column was used with a mobile phase containing acetonitrile, methanol, and 0.05% trifluoroacetic acid, and a gradient elution system with a 70-minute run time. Such parameters, particularly the proportion of organic modifiers and the column temperature (maintained at 40□°C), are known to significantly influence retention behaviour [43, 44]. The most relevant information which this HPLC run highlighted relates to this observed similarity in chromatographic profiles between encapsulated and unencapsulated Nisin in the present study. This confirms that there was no chemical degradation or peptide modification during encapsulation which could lead to shifts in retention time, peak shape deformation, or the appearance of degradation products. The absence of such changes implies that the ionic gelation method using alginate and chitosan is a mild encapsulation approach, preserving the structural integrity of Nisin and thus its activity which is critical for its antimicrobial efficacy. Similar stability has been observed in the discussed study by Bernela et al. [11] in which their HPLC results showed retention times consistent with unmodified Nisin, further confirming that encapsulation using biocompatible polymers does not adversely impact the active structure [11]. Furthermore, the HPLC results in this study complement the findings from FTIR spectroscopy, which showed no significant shifts in the functional group absorbances of Nisin after encapsulation. Together, these data affirm that both the primary structure and the functional groups of Nisin remained unaltered, which is crucial for maintaining its antimicrobial activity.

### 3.9. Morphology of Nisin 2A Microcapsules

In this study, the use of ionic gelation between alginate and calcium chloride allowed for the initial formation of a stable hydrogel pre-network. This was then further complexed with chitosan and Nisin through the electrostatic interactions between the negatively charged alginate moity and the positively charged chitosan and Nisin, resulting in a stable microcapsule. SEM imaging confirmed the morphological integrity of the synthesized microcapsules, which were mostly spherical or ellipsoidal with smooth surface textures and sizes ranging from 150–200 nm. While some particle aggregation was observed likely due to electrostatic interactions during drying or sample preparation, the overall morphology is consistent with similar chitosan-alginate systems such as seen in recent studies [45] with compact, smooth-surfaced particles with occasional clumping at a size of 200nm, highlighting effective encapsulation similar to those observed in the present study as shown in the representative micrograph (Figure 5). The size of the microcapsule was determined by measuring single occurring microcapsules from 10 image slices at the same magnification (50,000x) and calculating the average. However, aggregates with larger particle sizes can also be seen in the SEM micrograph, showing the propensity for the particles to aggregate.

**Figure 5.**
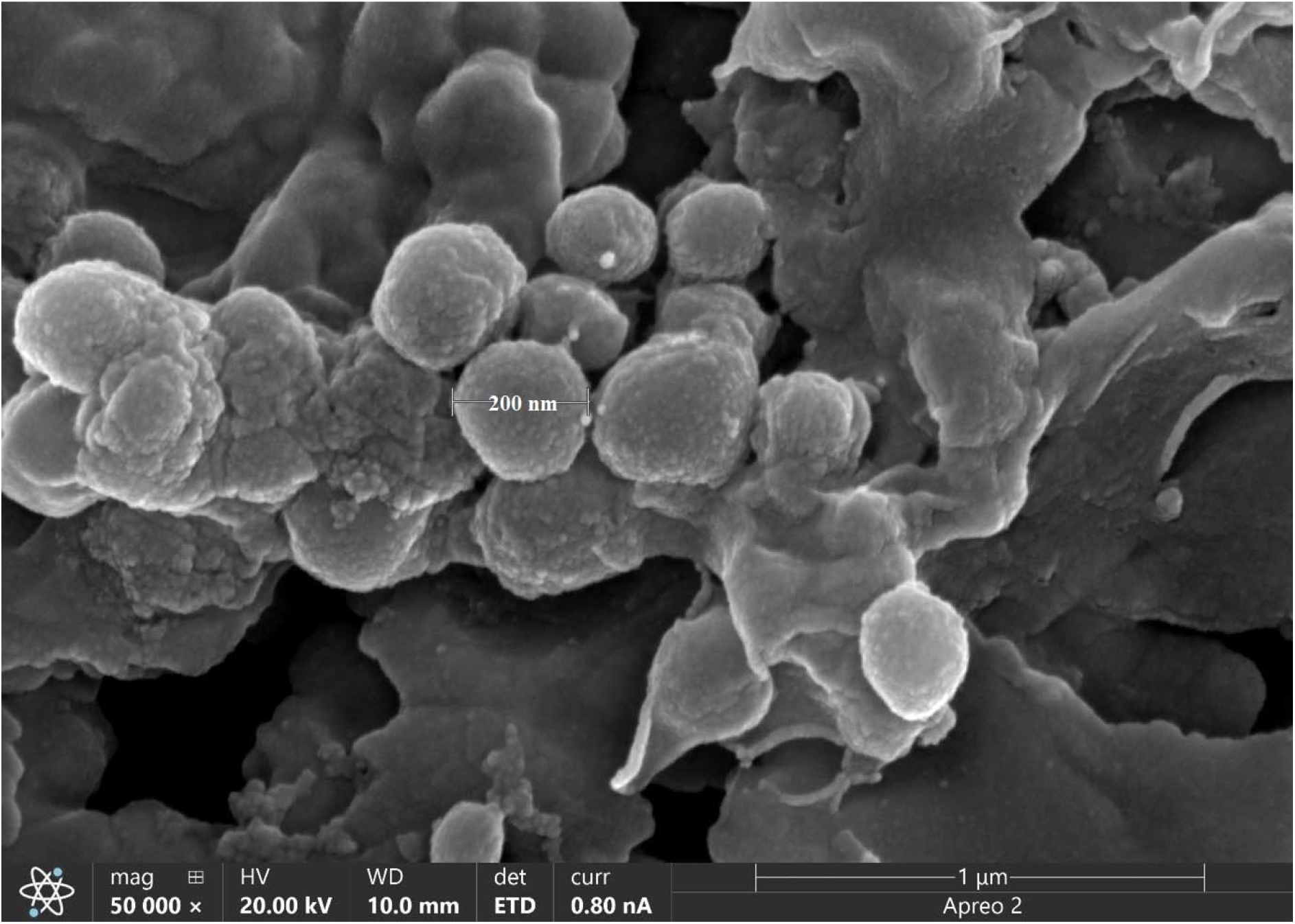
SEM micrograph of Nisin-chitosan-alginate microcapsules showing a size of 200nm

The morphology of microcapsules, particle shape, surface texture, and aggregation behaviour directly affect the applicability of the microcapsule in terms of encapsulation efficiency, release kinetics, and functionality in food systems or other areas of application [46–48]. Overall, the morphological features of the microcapsules obtained in this study are consistent with previously published studies using similar biopolymer systems and such morphology is typically associated with effective polymer–polymer interactions and stable encapsulation of hydrophilic bioactive compounds such as Nisin. For example, in a similar study by Hu et al. [49], it was demonstrated that Nisin encapsulated in chitosan nanoparticles exhibited spherical geometry and smooth external surfaces, with particle sizes of about 400nm, quite higher than the range reported in this present study. Their work, although with a lower encapsulation efficiency also highlighted that uniform morphology contributes to consistent antimicrobial release, as spherical microcapsules minimize surface defects that could lead to premature or uncontrolled diffusion of the active compound. The smooth and compact appearance of the microcapsules in this study suggests a well-formed polyelectrolyte complex between chitosan and alginate, resulting in a coherent shell that encapsulates Nisin efficiently. This structural integrity is particularly important for maintaining chemical stability and preventing leakage of Nisin during storage and application. This is further influenced by the strength of the linkage between chitosan and alginate which is determined by factors such as charge density and pH as shown in a related study by Aluani et al. [50] where a higher concentration and strength of sodium alginate results in smaller sized microcapsules with negative charge while higher concentration of chitosan resulted in larger capsules and positive charge, thus, directly affecting the formation of dense, spherical particles. Furthermore, the presence of aggregation of the microcapsules in some SEM fields indicates the tendency of particles to associate under certain conditions. This could be attributed to insufficient electrostatic repulsion during drying, as surface charge is lost in the absence of an aqueous medium. Similar phenomena have been reported in the literature [51] in which particle clustering in samples of alginate–chitosan capsules was observed and was attributed to high surface energy and collapse of the hydration shell. The occurrence of such aggregation does not necessarily reflect poor encapsulation quality but does suggest that modification of drying or storage protocols such as freeze-drying with cryoprotectants may be beneficial for maintaining dispersibility [52].

The size measurements obtained from SEM in this study correspond closely with those derived from DLS, supporting the reliability of the observed particle morphology and provides further insights into the colloidal stability and uniformity of the microcapsules. While SEM provides insight into the dry-state morphology, DLS assesses particles in suspension. The alignment between these two techniques confirms that the particles retain their size and shape in both solid and dispersed states, indicating structural stability.

To understand possible mechanism of bacteria inactivation by Nisin 2A, SEM of treated *Bacillus cereus* was undertaken. SEM revealed clear morphological disruptions in bacterial cells following treatment with Nisin 2A. Treated cells displayed a corrugated and irregular surface, along with visible leakage of intracellular contents (Figure 6a and 6b), in contrast to the smooth, undamaged morphology of untreated control cells. These findings are consistent with previous reports demonstrating membrane deformation and cytoplasmic leakage in *Listeria monocytogenes* and *Bacillus subtilis* upon exposure to lantibiotics such as Nisin and subtilin [53, 54]. However, while SEM provided qualitative evidence of membrane damage, its limited resolution prevented visualization of finer structural features such as pore formation. To overcome this would require the use of transmission electron microscopy (TEM) to enable high-resolution imaging of peptide-induced ultrastructural changes in bacterial cells, similar to the approach taken by Hartmann et al. for higher resolution of their samples [55]. However, this was not undertaken in this present study due to limitations in equipment availability.

**Figure 6.**
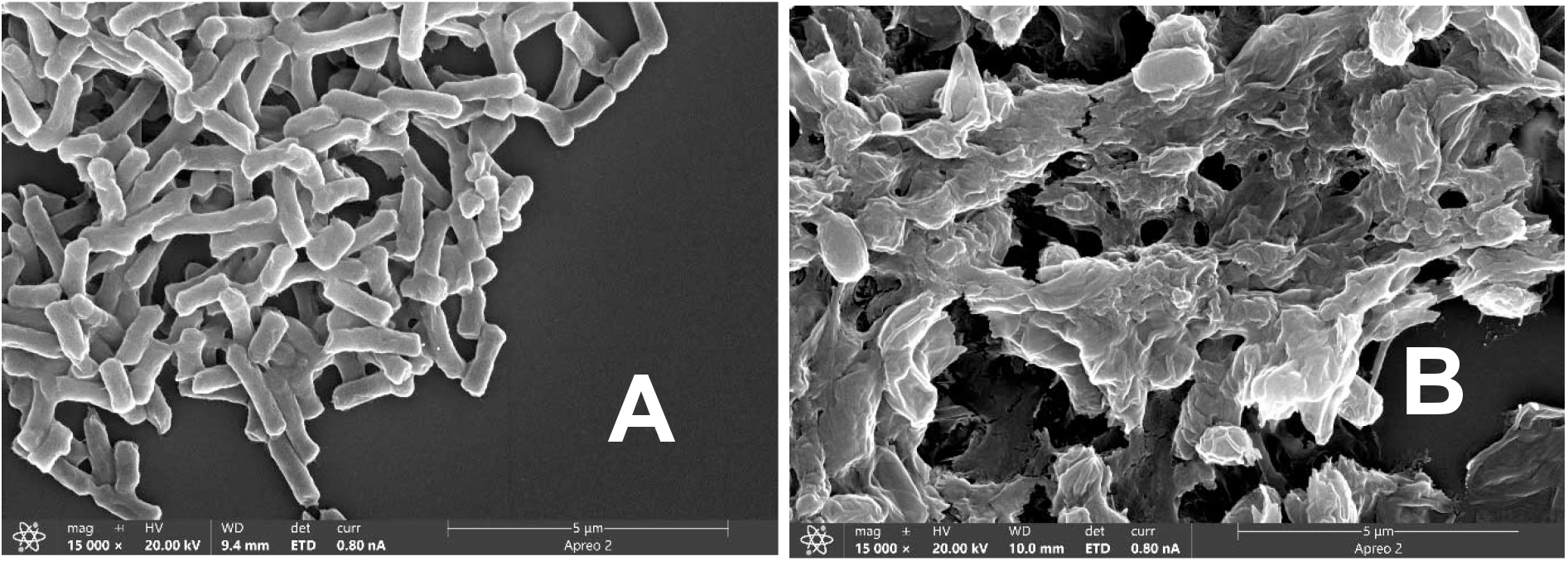
Scanning electron micrograph of *Bacillus cereus.* (A) untreated control, (B) Nisin 2A treated

### 3.10. FTIR analysis of Nisin-loaded microcapsules

The FTIR spectra of Nisin-loaded microcapsules were compared against spectra of pure Nisin, blank microcapsules, and the individual polymers to determine whether any structural modifications or chemical interactions occurred during the encapsulation process. Spectral analysis showed that the primary functional groups of Nisin including the OH, C=O, NH, and amide groups were preserved post-encapsulation, indicating that no significant chemical interactions occurred between Nisin and the biopolymer matrix comprising chitosan and alginate. This observation aligns with previous findings in similar encapsulation systems including those of Bernela et al. [11] and Hu et al. [56] and confirms that the encapsulation process maintains the structural integrity and bioactivity of Nisin. FTIR analysis reveals a spectrum of Nisin at 3288 cm^−1^, possibly due to OH stretching of COOH group, and another peak at 2960 cm^−1^ due to C–H Stretching, while the peak at 1527 cm^−1^ is due to bending of primary amines. In addition, the peak obtained at 1232 cm^−1^ results from the O–H group while that at 1645 cm^−1^ is due to amide groups. Analysis of the spectra of Nisin-loaded microcapsules shows a peak at 3284 cm^−1^ due to free O–H groups of COOH; the peak at 2935 cm^−1^ results from C–H stretching, and the peak at 1085 cm^−1^ correlates with secondary hydroxyl groups. The predominant functional groups of the polymeric material within the surface of the microcapsule have chemical characteristics similar to those of Nisin. Thus, this spectral analysis indicates that chemical interactions between functional groups of Nisin and the polymeric substances which can lead to the alteration of the chemical structure of Nisin did not occur. Therefore, Nisin can remain stable within the microcapsule. This is similar to results obtained in a similar study which encapsulated alpha-1 antitrypsin in poly (D, L lactide-co glycolide) (PLGA) nanoparticles, with no interaction observed in FTIR spectral results [57], and in another study which encapsulated Nisin in an alginate-chitosan-pluronic composite microcapsule [11]. Similarly, the FTIR spectrum of Nisin observed in similar studies [56] showed the absence of significant shifts in the peaks of functional groups (1645 to 1634 cm□¹ for the amide band for instance) suggesting that no chemical interactions occurred between Nisin and the encapsulating polymers that could alter the structure of Nisin, which corroborates the findings made in this present study that encapsulation effectively preserved the chemical stability of Nisin. A related observation in this study is the similarity in chemical profiles between blank microcapsules and the Nisin-loaded ones, further suggesting that Nisin was incorporated primarily through physical entrapment mechanisms, such as hydrogen bonding and ionic interactions within the capsules, without altering its intrinsic structure. These results align with studies on other protein or peptide encapsulation systems, such as in the PLGA nanoparticles study by Pirooznia et al. [57] which showed through FTIR spectra that no chemical alterations occurred during the encapsulation process. Taken together, the FTIR data in this study confirm that Nisin remains structurally intact after encapsulation, and that the chitosan-alginate matrix provides a chemically inert but physically compatible environment. This is essential for preserving the bioactivity and functional efficacy of Nisin in food preservation applications. Moreover, the maintenance of Nisin’s characteristic functional groups underlines the effectiveness of the ionic gelation method as a gentle and non-denaturing encapsulation technique. Figure 7 shows FTIR spectra of Nisin, blank microcapsules and Nisin-loaded microcapsules.

**Figure 7.**
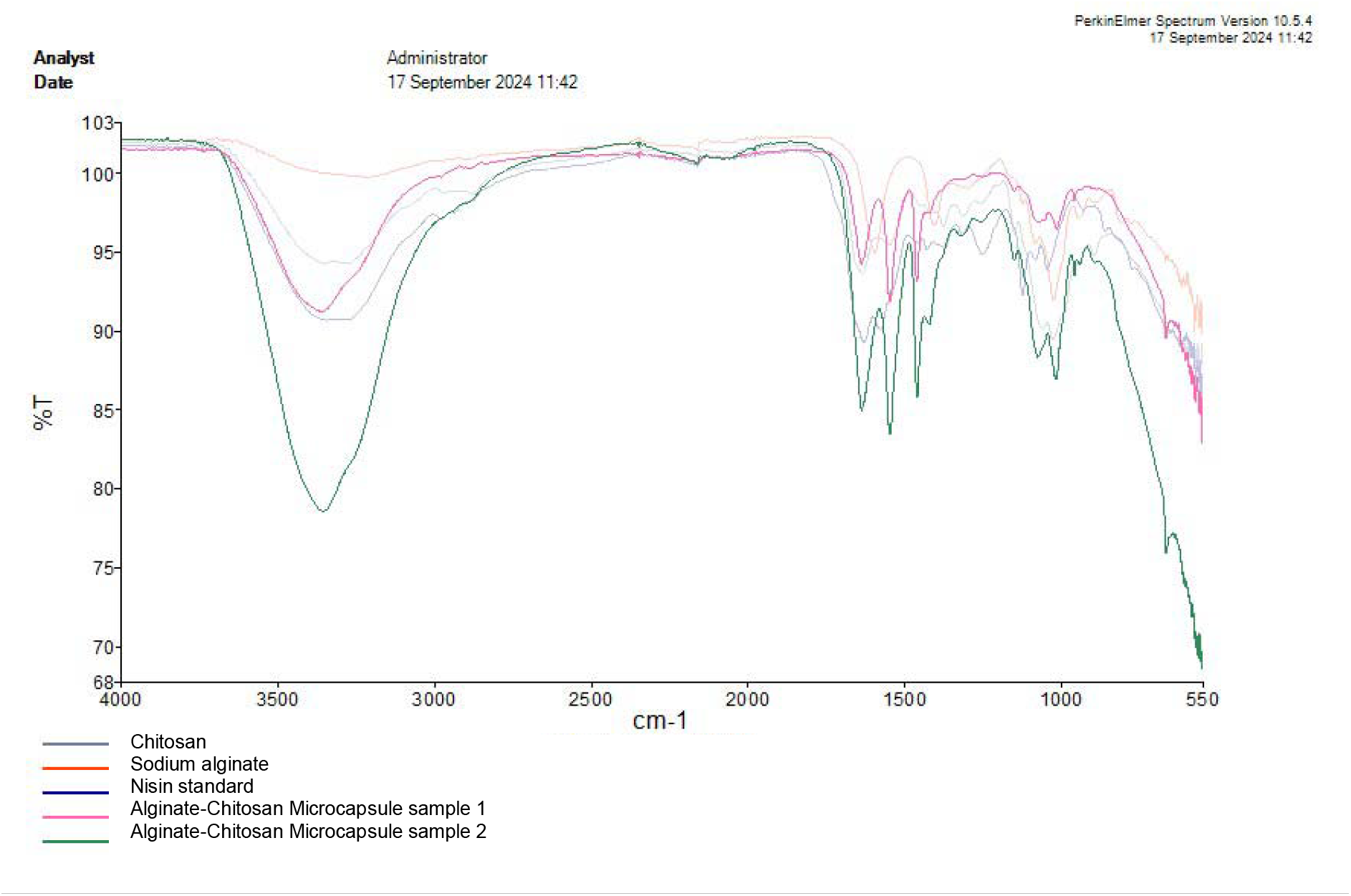
FTIR spectra of Nisin-loaded microcapsules

### 3.11. Evaluation of the stability of Nisin-chitosan-alginate microcapsules through 21 days of cold storage at 4°C

To validate the biological function of the encapsulated product, the antimicrobial activity of the Nisin-loaded microcapsules was evaluated against *Bacillus cereus* following a 21-day refrigerated storage period using 0.5 mg/L of the Nisin-chitosan-alginate microcapsule. The results showed that Nisin-loaded microcapsules retained robust antimicrobial activity against *Bacillus cereus*, over the 21-day period at 4□°C, as evidenced by similar zones of inhibition with diameter of about 10mm obtained in control samples across all days of storage, suggesting that the encapsulation process did not diminish the bioactivity of Nisin. This stability contrasts with the behaviour of unencapsulated Nisin, which is often prone to deactivation during prolonged storage. These findings align with previous research demonstrating that Nisin maintains its antimicrobial efficacy when encapsulated in biopolymer systems, particularly under refrigerated storage conditions. For example, a similar study by Wienke et al. [58] investigating polyelectrolyte-coated liposomes as nanostructured systems for the delivery of Nisin found that the liposomes that contained Nisin demonstrated an enhanced antimicrobial activity (from 3200 to 6400 AU/mL) against *B. cereus* when Nisin was encapsulated with chitosan or maltodextrin coatings. They concluded that while the antimicrobial activity of unencapsulated Nisin is usually comparable to that of encapsulated Nisin as shown in this present study, the unencapsulated Nisin typically performs poorly when applied directly to food as they may interact with proteins and fats which reduce their level of antimicrobial effectiveness. While their findings emphasized the synergistic effect of polyelectrolyte coatings, this present study provides a simpler but equally effective encapsulation alternative using a chitosan-alginate system, showing that similar preservation of antimicrobial function can be achieved without liposomal carriers.

Importantly, this study demonstrated that storage time did not significantly affect the antimicrobial performance of encapsulated Nisin under refrigerated storage. It has been previously demonstrated that during prolonged storage, unencapsulated Nisin tends to bind to proteins, lipids, or polyphenols in food, which significantly reduces its bioavailability and antimicrobial effect [59–61]. In contrast, encapsulation provides a physical barrier that protects Nisin from premature interactions and degradation, leading to prolonged antimicrobial activity in complex environments such as fruit juices, dairy, or meat products. A previously cited study by Bernela et al. [11] supports the conclusion that chitosan-based delivery systems not only protect the antimicrobial properties but may also enhance its efficacy through synergistic effects. Chitosan itself exhibits mild antimicrobial activity due to its polycationic nature and interaction with bacterial membranes, which may complement Nisin’s pore-forming mechanism. The combined antimicrobial action likely contributes to the consistency in performance observed across all storage intervals in our study, making such a system suitable for long-term preservation strategies in cold-stored food products.

### 3.12. *In vitro* release of Nisin from Nisin-chitosan-alginate microcapsules

The release kinetics of the encapsulated Nisin is an important parameter for the development of an effective and reliable food preservation system. In this study, the *in vitro* release profile of Nisin from chitosan-alginate microcapsules was assessed under controlled conditions using phosphate buffer (pH 5.0) over a 10-day period and revealed a sustained and controlled biphasic release, with approximately 40% of Nisin released within the first five hours following incubation/storage, reaching 60% by 24 hours, and up to 86% by 240 hours (10 days) (Figure 8). These results indicate the successful formation of a polymer matrix capable of modulating Nisin release, thus ensuring the sustained presence of the antimicrobial agent over a longer period as compared to the rapid release observed in unencapsulated controls, where more than 80% of free Nisin diffused within the first 2 hours of incubation/storage. This biphasic release pattern which comprises an initial moderate burst followed by sustained release is characteristic of polyelectrolyte complexes such as chitosan-alginate complex which provides a physical entrapment and electrostatic interactions with Nisin, thus ensuring a slower rate of diffusion [8]. The initial release observed in this study can be attributed to surface-associated Nisin within the chitosan-alginate matrix or superficial desorption of loosely bound Nisin, while the slower, prolonged release phase results from Nisin diffusing through the denser internal matrix of the microcapsules/interpolyelectrolyte network formed between chitosan and alginate, which provides a semi-permeable barrier around the Nisin molecules [8, 62]. Such biphasic release kinetics have been frequently reported in polymeric encapsulation systems, such as in the earlier study by Zohri et al. [9] where a similar release trend in Nisin-alginate particles, in a pH-dependent manner was described.

**Figure 8.**
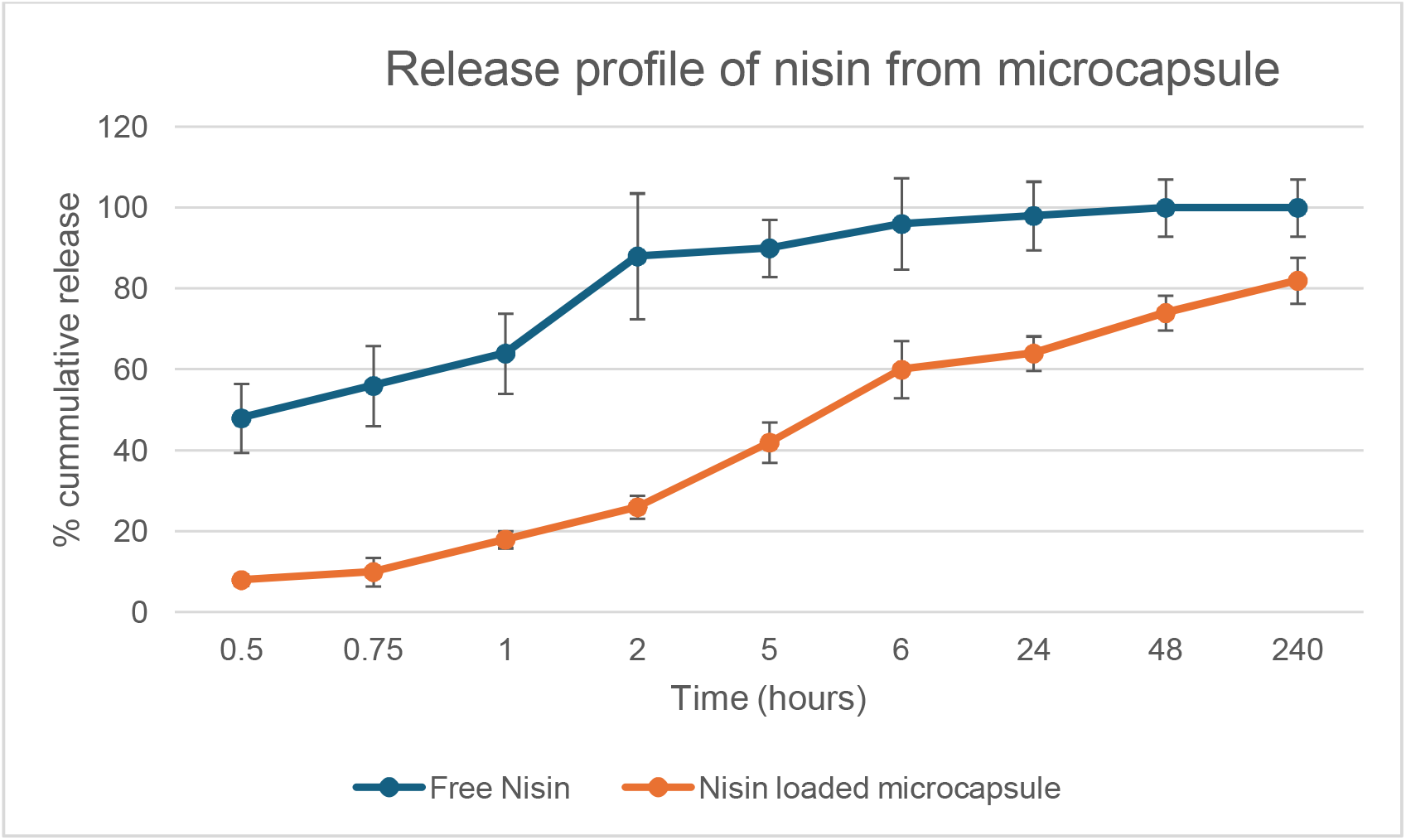
Release profile of Nisin from chitosan-alginate microcapsule

A lower pH environment can slow Nisin release from microcapsules due to stronger electrostatic interactions between the cationic peptide and the anionic polymers [9]. In this present study, release was optimized at a mildly acidic pH (pH 5.0), which mimics the natural environment of many acidic foods (e.g., fruit juices). At this pH, chitosan is partially protonated, contributing to matrix stability, while Nisin retains its antimicrobial structure. The results obtained from this present study highlights this pH-dependent release of Nisin, with optimal release kinetics obtained at a pH of 5.0. This is similar to that obtained by Fael et al.’s study on Nisin/polyanion layer-by-layer films [63] where the stability of Nisin/polyanion films at three different pH values were investigated and it was found that Nisin release from polyacrylic acid films followed a biphasic release process dependent on pH, with a burst release of about 85% within the first hour and subsequent slower release and disintegration in 5-24 hours at pH of 9.5 whereas less than 60% was released within the same time period from phosphate buffered solution at pH 6.0 and 7.4. This observation was attributed to the less net positive charge on Nisin molecule with increasing pH and groups in poly acrylic acid. This was attributed to reduced electrostatic interactions at higher pH due to the loss of Nisin’s positive charge and greater swelling of the polymetric matrix. The results obtained in this present study similarly suggest that Nisin’s net charge and the state of ionization of alginate and chitosan significantly influence diffusion and release behaviour, and this is more stable at the slightly acidic to neutral pH favoured in this study which is more relevant to food systems. In this study, the controlled release behaviour observed is functionally advantageous because in food systems, a slow and sustained release ensures long-term antimicrobial protection, prevents the rapid depletion of the bioactive compound, thus extending the shelf life of the product [64]. Encapsulation helps to achieve this protection within the matrix and releases gradually in response to environmental conditions, unlike free Nisin which can be rapidly inactivated in the food matrix through interactions with food proteins, lipids, or enzymes. This provides a more effective strategy for microbial control, particularly against resistant spore-forming bacteria such as *Bacillus cereus* or *Alicyclobacillus* spp., which are common spoilage agents in acidic beverages like apple juice, as the encapsulated system described here shows promise for long-term delivery and consistent antimicrobial action over time. Importantly, the extended release of Nisin may help in maintaining sub-lethal concentrations of the antimicrobial over prolonged periods (above 10 days in this present study), which can inhibit microbial growth without inducing resistance. This is especially important for targeting spore-forming bacteria like *Bacillus cereus*, which require sustained exposure to antimicrobials for effective inhibition of germination and spore outgrowth phases. The findings from this release affirms the validity of the encapsulation strategy and supports its potential utility in commercial food systems that demand both safety and shelf-life extension. Overall, the *in vitro* release results obtained in this study demonstrates that Nisin-chitosan-alginate microcapsules offer a stable and pH-responsive delivery system for Nisin, capable of modulating its release over several days. The ability to sustain release under conditions relevant to acidic foods makes this system particularly attractive for food preservation applications especially in the juice sector.

## CONCLUSION

This study successfully demonstrated the microencapsulation of Nisin using a chitosan-alginate biopolymer matrix via ionic gelation and polyelectrolyte complexation, to enhance the antimicrobial peptide’s stability and efficacy for potential application in food preservation, particularly against spoilage organisms such as *Bacillus cereus*. The encapsulation protocol was optimized by controlling polymer ratios, calcium ion concentration, and pH and resulted in the formation of spherical, nanoscale microcapsules (150–200 nm) with smooth surface morphology as confirmed by SEM. Dynamic Light Scattering (DLS) and Zeta potential analysis showed that the microcapsules were monodisperse (PDI ∼0.302) and electrostatically stable (+36.4 mV), supporting their long-term colloidal stability in aqueous systems. These stability assessments were conducted under controlled laboratory conditions, providing a basis for further validation in complex food matrices. Furthermore, the study highlighted the critical role of pH in modulating polymer ionization and interpolymer interactions, which in turn influenced microcapsule formation, stability, and encapsulation efficiency. Encapsulation efficiency reached 65%, confirming effective retention of Nisin within the polymer matrix under optimized conditions. This encapsulation efficiency was closely associated with the controlled pH and polymer ratio conditions employed during microcapsule formation. Importantly, FTIR spectroscopy and HPLC analysis demonstrated that no significant chemical interaction or structural degradation of Nisin occurred during the encapsulation process, indicating that the method preserved the molecular integrity of the active compound. Functionally, the antimicrobial activity of the encapsulated Nisin remained consistent over 21 days of refrigerated storage, exhibiting inhibition zones similar to those of commercial Nisin (Ultrapure Nisin) when tested against *B. cereus*. Overall, this research provides strong evidence that chitosan-alginate microencapsulation is a viable, non-denaturing method for delivering and stabilizing Nisin in food systems. The use of GRAS-status biopolymers further supports the regulatory compatibility of this encapsulation system for food applications. The resulting microcapsules demonstrate promising potential for use in the natural preservation of acidic beverages such as apple juice, with relevance for broader food safety and shelf-life enhancement applications. The comprehensive physicochemical characterization and bioactivity assessments undertaken in this study confirms the functional integrity, structural stability, and practical applicability of Nisin-loaded chitosan-alginate microcapsules. Further work in real food matrices and scalability assessments, alongside continued optimization of formulation parameters and scale-up validation, will be necessary to translate this effective GRAS-based encapsulation system for Nisin into its full commercial potential in the food industry.

## Author Contributions

Conceptualization, C.K.A.; Methodology, H.O., C.K.A.; Writing—original draft, H.O. and C.K.A.; Writing—review and editing, C.K.A., H.O. and T.M.; Validation, T.M.; Supervision, H.O. All authors were involved in the writing and editing of this manuscript and have approved its submission for publication. All authors have read and agreed to the published version of the manuscript.

## Funding

This research received no external funding.

## Institutional Review Board Statement

Not applicable.

## Informed Consent Statement

Not applicable.

## Data Availability Statement

No new data were created or analyzed in this study. Data sharing is not applicable to this article.

## Conflicts of Interest

The authors declare no conflict of interest.

## Notes

### Competing Interest Statement

The authors have declared no competing interest.

